# Essential functions of MLL1 and MLL2 in retinal development and cone cell maintenance

**DOI:** 10.1101/2021.11.15.468679

**Authors:** Chi Sun, Xiaodong Zhang, Philip A. Ruzycki, Shiming Chen

## Abstract

MLL1 (KMT2A) and MLL2 (KMT2B) are homologous members of the mixed-lineage leukemia (MLL) family of histone methyltransferases involved in epigenomic transcriptional regulation. Their sequence variants have been associated with neurological and psychological disorders, but little is known about their roles and mechanism of action in CNS development. Using mouse retina as a model, we previously reported MLL1’s role in retinal neurogenesis and horizontal cell maintenance. Here we determine roles of MLL2 and MLL1/MLL2 together in retinal development using conditional knockout (CKO) mice. Deleting *Mll2* from Chx10*+* retinal progenitors resulted in a similar phenotype as *Mll1 CKO*, but removal of both alleles produced much more severe deficits than each single *CKO*: 1-month *double CKO* mutants displayed null light responses in electroretinogram; thin retinal layers, including shorter photoreceptor outer segments with impaired phototransduction gene expression; and reduced numbers of M-cones, horizontal and amacrine neurons, followed by fast retinal degeneration. Despite moderately reduced progenitor cell proliferation at P0, the neurogenic capacity was largely maintained in *double CKO* mutants. However, upregulated apoptosis and reactive gliosis were detected during postnatal retinal development. Finally, the removal of both MLLs in fated rods produced a normal phenotype, but the CKO in M-cones impaired M-cone function and survival, indicating both cell non-autonomous and autonomous mechanisms. Altogether, our results suggest that MLL1/MLL2 play redundant roles in maintaining specific retinal neurons after cell fate specification and are essential for establishing functional neural networks.

## INTRODUCTION

The mixed-lineage leukemia (MLL) family of histone lysine methyltransferases (KMTs) catalyze the methylation at lysine (K) 4 of histone H3 (Crump & Milne, 2019; Shilatifard, 2008). Their products, mono-, di- and tri-methylation of H3K4 are considered activating histone marks that are found at promoters of transcribed genes (Eissenberg & Shilatifard, 2010; Gu & Lee, 2013). The MLL family has six members, namely, MLL1 (KMT2A), MLL2 (KMT2B), MLL3 (KMT2C), MLL4 (KMT2D), SET1DA and SET1DB, all of which contain a conserved catalytic SET domain (Allis et al., 2007; Crump & Milne, 2019). These MLL enzymes usually form large multi-protein complexes (Dou et al., 2006; Shinsky et al., 2015; Yokoyama et al., 2004) that mediate chromatin remodeling and transcriptional regulation during tissue genesis (Cenik & Shilatifard, 2021; Glaser et al., 2006; Krivtsov & Armstrong, 2007; Slany, 2016). MLL1/MLL2 form a distinct macromolecular complex (Crump & Milne, 2019; Sze & Shilatifard, 2016) found in many cell types of the central nervous system (CNS) (Brightman et al., 2018; Jakovcevski et al., 2015; Kerimoglu et al., 2017). Human mutations in MLL1 (KMT2A) and MLL2 (KMT2B) have been associated with cancer and syndromic neurological and psychiatric disorders, such as dystonia, intellectual disability and autism with variable age onsets (Jones et al., 2012; Meyers et al., 2017). However, their functions and mechanisms of action in the CNS development and neuronal disease pathogenesis remain to be elucidated.

The retina is the tissue of the CNS specialized for the vision sensation. It consists of conserved and well-characterized neuronal cell types, making it an ideal model for studying MLL functions in neurogenesis, neural network formation and maintenance. Retinal cells are organized into a multi-laminar structure in a mature retina (Bassett & Wallace, 2012; Cepko, 2014; Cepko et al., 1996; Young, 1985). The outer nuclear layer (ONL) contains rod and cone photoreceptor cells, where the light signal is converted into the neuronal signal. The inner nuclear layer (INL) hosts neurons of horizontal (HC), bipolar (BC) and amacrine cells (AC), as well as Müller glia (MG). The ganglion cell layer (GCL) contains retinal ganglion cells (GC) innervating to the brain. Synaptic connections between these layers form two additional layers known as the inner (IPL) and outer plexiform layers (OPL). Cell-type specification of the retina is largely conserved across vertebrate species (Austin & Cepko, 1995; Livesey & Cepko, 2001; Lu et al., 2020; Stenkamp, 2015; Wu et al., 2018). In mice, GC, cone photoreceptors and HC are fated in the first wave of cell birth during prenatal retinal neurogenesis, followed by the later birth waves of AC, rod photoreceptors, BC and MG during postnatal days (P) 0 to P5 (Cepko, 2014; Emerson et al., 2013; Hafler et al., 2012; Turner et al., 1990; Young, 1985). After cell fate specification, neuronal differentiation and synaptogenesis begin to establish connections between rod/cone primary neurons and secondary/tertiary neurons. By the end of three weeks, a functional neural network is established to mediate visual signal transmission and processing (Blanks et al., 1974; Furukawa et al., 2020; Guido, 2018; Soto et al., 2012; Szatko et al., 2020; Tian, 2004).

Retinal cell fate specification and differentiation are regulated by a network of transcription factors (TF) (Hennig et al., 2008; Mellough et al., 2019; Swaroop et al., 2010). CHX10 (also known as VSX2) is one of the earliest expressed TFs in the presumptive neural retina, and is required for retinal progenitor cell (RPC) proliferation and neurogenesis (Burmeister et al., 1996; Green et al., 2003; Liu et al., 1994; Livne-bar et al., 2006). CHX10 is also required for bipolar cell specification at a later development stage (Kim et al., 2008; Morrow et al., 2008). OTX2 is essential for establishing the photoreceptor lineage (Koike et al., 2007; Nishida et al., 2003) by inducing the expression of CRX, an OTX-like TF required for photoreceptor development and maintenance (Chen et al., 1997; Furukawa et al., 1997; Hennig et al., 2008). TFs acting downstream of OTX2/CRX govern photoreceptor specification, including NRL (Mears et al., 2001) for rods, THRB2 (Roberts et al., 2006) and RXRG (Roberts et al., 2005) for cones, specifically in mouse, for short-wavelength (S) and medium-wavelength (M) cones. TFs required for the development of INL and GCL neurons include Onecut1 (OC1) and PROX1 for horizontal cells (Dyer et al., 2003; Wu et al., 2013), PAX6 for amacrine cells (Remez et al., 2017), and POU4F1 (BRN3A) for ganglion cells (Nadal-Nicolás et al., 2009).

In order to express specific genes in a given cell type, the chromatin at these gene loci must be configured to allow access by the transcription machinery. Specific TFs can regulate local chromatin remodeling during the retinal development by recruiting co-activators and histone modification enzymes at target genes (Li et al., 2007; Zhang et al., 2015). For example, CRX is required for the postnatal developing photoreceptors to gain DNA accessibility and write active histone marks H3K4me3 and H3K27Ac at photoreceptor gene loci (Ruzycki et al., 2018), which is important for the transcription cascade and visual function.

We previously reported, using conditional knockout mice, that MLL1 plays essential roles in regulating retinal progenitor cell proliferation, maintenance of horizontal neurons and formation of functional synapses between photoreceptors and inner neurons (Brightman et al., 2018). Here we aimed to unveil the roles of MLL2 and MLL1/MLL2 together in the development and maintenance of retinal neurons by characterizing the phenotypes of retina-specific knockout of *Mll2* and both *Mll1* and *Mll2*. We found that during the functional development, two MLLs played redundant roles in maintaining the retinal structure and function of photoreceptors and three types of inner retinal neurons. Both cell autonomous and non-autonomous functions contributed to cone/rod maintenance. These results implicated complex mechanisms of action of MLL1 and MLL2 in the retinal development and maintenance.

## RESULTS

### Generation of *Mll1* and *Mll2* conditional knockouts in mouse retinae

Due to the embryonic lethality of *Mll1 or Mll2* deficiency, we utilized *Cre-LoxP*-mediated conditional knockout (CKO) to eliminate the function of each gene individually or simultaneously in selected retinal cell types. Cre excision produced a non-functional MLL1 protein lacking the nuclear targeting sequences (NTS) (Gan et al., 2010) and/or a short non-functional MLL2 peptide resulted from a frameshift at exon 2 (Glaser et al., 2006) (Figure 1A). To investigate the roles of MLL1 and MLL2 in retinal neurogenesis, we crossed *Mll1^f/f^* and/or *Mll2^f/f^* mice to *Chx10-Cre-iresGFP* (Rowan & Cepko, 2004) mice, which expressed Cre recombinase in retinal progenitor cells. To confirm the success of Cre-mediated excision, we performed quantitative RT-PCR (qRT-PCR) analysis to determine *Mll1* and *Mll2* transcript levels in the mutant retinae at postnatal day P0, P14, and 1 month (1MO) (Figure 1B-D). Detectable transcript levels of *Mll1* and *Mll2* were significantly reduced in *Chx10Cre*^+^*Mll1^f/f^Mll2^f/f^* (*double CKO*) retinae at all tested ages as compared to *Chx10Cre^-^* (*CreNeg*) samples with the lowest level seen at 1MO. Notably, at all tested ages, reduction of *Mll1* or *Mll2* transcript levels in the *double CKO* retinae was comparable to the corresponding *single CKO*. Furthermore, in either case of *single CKO* retinae, no increase in the transcript level of the counterpart *Mll* gene was detected, suggesting no compensatory effect between *Mll1* and *Mll2* expression in *single CKO* retinae.

**Figure 1.**
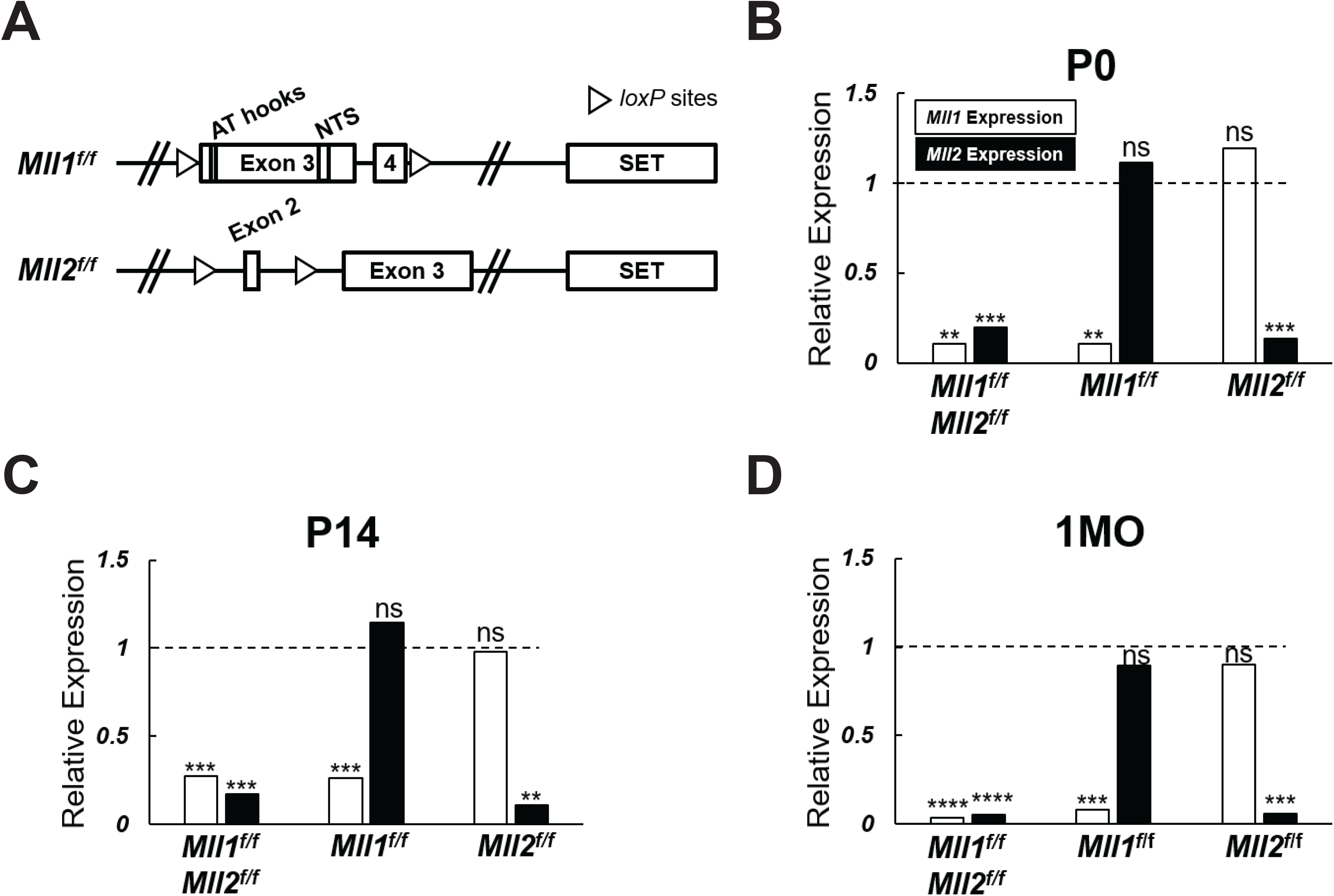
*Mll1* and *Mll2* expression in conditional-knockout retinae. (**A**) Diagrams of wide-type *Mll1* and *Mll2* genes indicating the positions of SET domains, motifs and *loxP* sites. Cre recombinase-mediated *Mll1* knockout removes exons 3 and 4 which encode the motifs of AT hooks and nuclear targeting signals (NTS) (Gan et al., 2010). Cre recombinase-mediated *Mll2* knockout removes exon 2 and invokes a frameshift in the mRNA (Glaser et al., 2006). (**B-D**) qRT-PCR analysis of *Mll1* and *Mll2* expression in P0, P14 and 1 month-old (1MO) mutant retinae. Results are plotted as relative expression to *Chx10Cre^-^* littermate controls (n≥4). Asterisks (**, ***, ****) denote *p* ≤ 0.01, *p* ≤ 0.001, *p* ≤ 0.0001 respectively by one-way ANOVA with Tukey’s multiple comparisons, and ns means not significant.

### *Mll1* and *Mll2* deficiency perturbs retinal function more severely than each single *CKO*

We performed electroretinography (ERG) to determine the effects of *Mll1* and *Mll2* deficiency on the retinal function. 1MO *double CKO* mutants showed no ERG responses in both dark-adapted (Figure 2A-B, purple line) and light-adapted (Figure 2C, purple line) conditions, indicating the absence of both rod- and cone-driven retinal function. In contrast, 1MO *single CKO* mutants produced detectable ERG responses (Figure 2A-C, red and green lines), but had significantly decreased A and B wave amplitudes at high light intensities as compared to those of *CreNeg* littermates. The ERG reductions were more pronounced in *Chx10Cre*^+^*Mll1^f/f^* than in *Chx10Cre*^+^*Mll2^f/f^* (Figure 2A-C, green line vs red line). Similar patterns of ERG changes were also noticeable at 2MO (Supplemental Figure 1A). These results provided the first piece of evidence for distinct and redundant roles of MLL1/MLL2 in the development and maintenance of retinal function. Compound heterozygous mutants *Chx10Cre*^+^*Mll1^WT/f^Mll2^f/f^* and *Chx10Cre*^+^*Mll1^f/f^Mll2^WT/f^* showed similar ERG defects as those of the *single CKO* mutants (Supplemental Figure 1B-C) at 1MO and 2MO, indicating that at least one copy of *Mll1* or *Mll2* allele was essential for retinal function.

**Figure 2.**
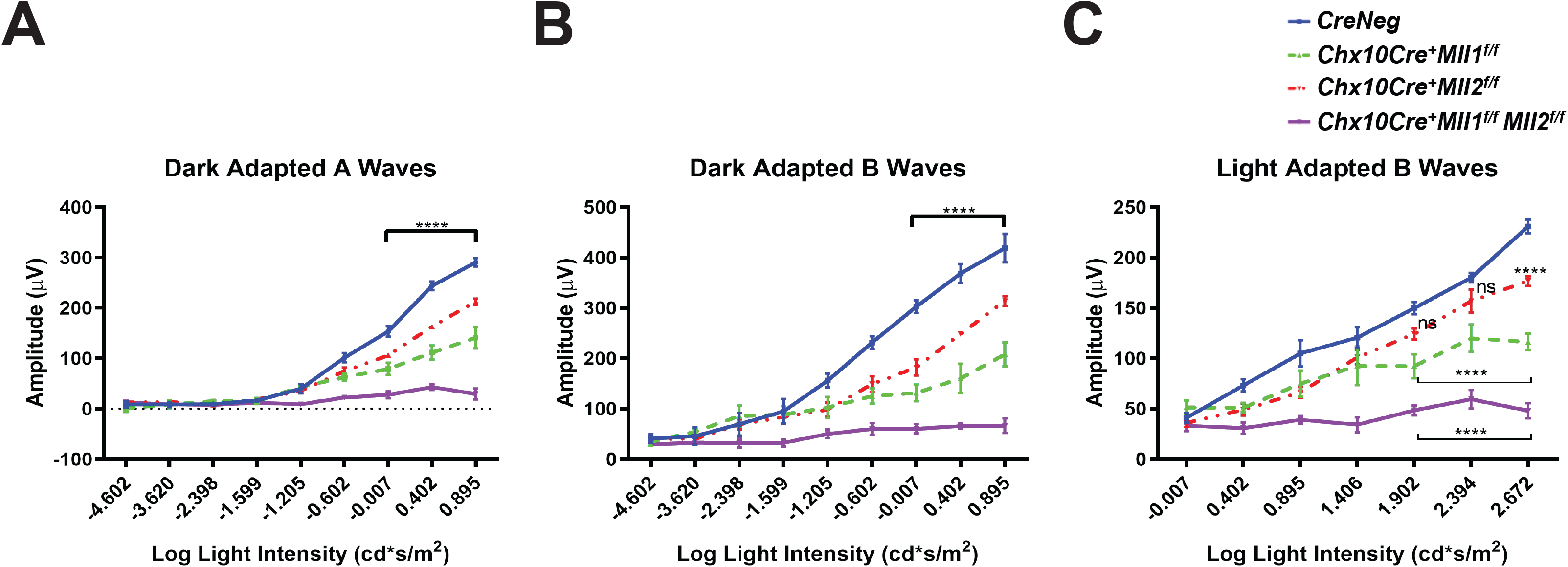
Impaired retinal function of *Mll* mutants at 1MO. (**A**) Dark adapted A-wave electroretinogram (ERG) analysis of 1MO mice of the indicated genotypes. Mean amplitudes (µV) are plotted against stimulus light intensity. Error bars represent SEM (n ≥ 4). (**B**) Dark adapted B-wave ERG analysis of 1MO mutant and control mice (n ≥ 4). (**C**) Light adapted B-wave ERG analysis of 1MO mutant and control mice (n ≥ 4). All statistics is done by two-way ANOVA and Tukey’s multiple comparisons. Asterisks (****) denote *p* ≤ 0.0001, and ns means not significant.

### *Mll1* and *Mll2* double *CKO* mice develop thinner retinae and undergo fast retinal degeneration in young adults

To determine if the ERG defects in the mutant retinae were indicative of changes in the retinal structure and cellular morphology, we firstly examined the retinal cross-sections by hematoxylin-and-eosin (H&E) staining. At 1MO, all mutants displayed well-laminated neurons (Figure 3A). However, as compared to the *CreNeg* control, the *double CKO* retinae had significant reductions in overall retinal thickness, including ONL (Figure 3B), INL (Figure 3C), and outer segments (OS) of the photoreceptors (Figure 3D). Mislocalized cells were found (indicated by asterisks in Figure 3A) within the OS and IPL of the *double CKO* mutant. At 2MO, *single CKO* mice retained the laminated retinal morphology, despite a few mislocalized cells found at the IPL (Figure 3A, bottom left panels). In contrast, the *double CKO* retinae became much thinner than that at 1MO, suggesting severe degeneration and cell death (Figure 3A). By 3MO, both ONL and INL of the *double CKO* retinae degenerated further with only 1-2 rows of cells remaining in each layer (Figure 3A). Also, the OS length in the *double CKO* retinae was further reduced at 2MO and 3MO. Thus, loss of both *Mll1* and *Mll2* resulted in failure of morphological maintenance of the retinae, particularly affecting neuronal survival in the ONL and INL. Interestingly, 2MO compound heterozygous mutants, *Chx10Cre*^+^*Mll1^WT/f^Mll2^f/f^* and *Chx10Cre*^+^*Mll1^f/f^Mll2^WT/f^* appeared less affected in retinal morphology as compared to the double mutants (Supplemental Figure 2A), again suggesting at least one copy of either *Mll1* or *Mll2* allele was required for the maintenance of retinal integrity.

**Figure 3.**
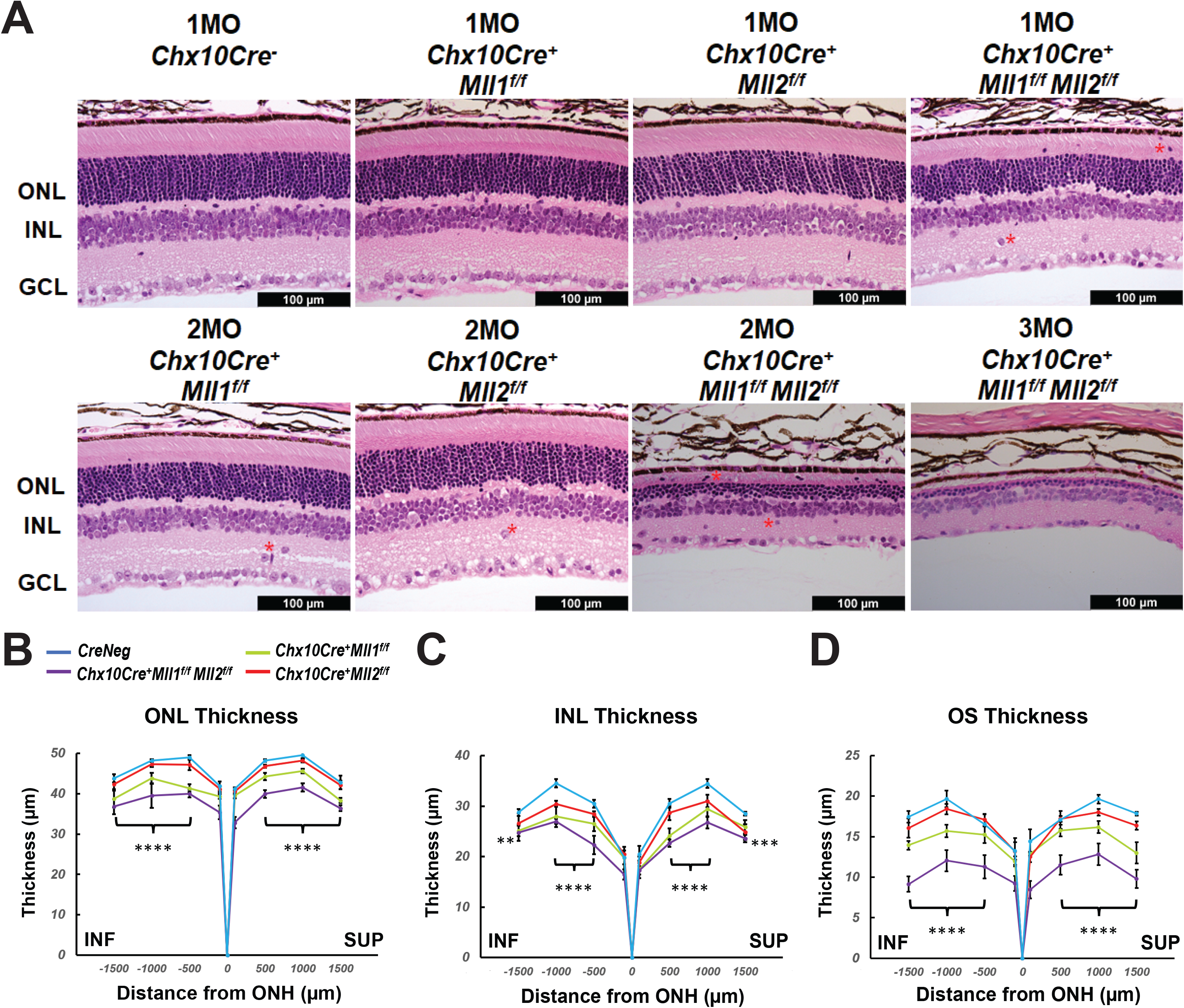
Morphological defects of *Mll* mutants. (**A**) Hematoxylin and Eosin (H&E) cross-section staining of mature mutant and control retinae. Top panels show representative images of 1MO retinae of the indicated genotypes. Bottom panels show representative images of 2MO or 3MO retinae of the indicated genotypes. Red asterisks indicate mislocalized cells. ONL: outer nuclear layer; INL: inner nuclear layer; GCL: ganglion cell layer. Scale bar = 100µm for all image panels. (**B**) ONL thickness (µm) in 1MO retinae at the indicated positions from the optic nerve head (ONH). SUP and INF indicate superior and inferior sides of the retina. Error bars represent SEM (n≥4). (**C**) INL thickness in 1MO mutant and control retinae (n≥4). (**D**) Outer segment (OS) thickness in 1MO mutant and control retinae (n≥4). All statistics is done by two-way ANOVA and Tukey’s multiple comparisons. Results of statistical analysis between the *double CKO* and *CreNeg* retinae are shown. Asterisks (**, ***, ****) denote *p* ≤ 0.01, *p* ≤ 0.001, *p* ≤ 0.0001 respectively.

We next examined changes in the retinal morphology at early ages of postnatal development. No morphological or thickness abnormalities were found in the new-born P0 *double CKO* retinae (Supplemental Figure 3A, B). However, decreased ONL and INL thickness was found in P14 *double CKO* retinae as compared to *CreNeg* samples (Supplemental Figure 3C, D). These results suggested that *Mll1* and *Mll2* deficiency interfered with maintenance but not genesis of retinal neurons.

### *Mll1* and *Mll2* deficiency induces progressive degeneration of M-cones, horizontal and amacrine cells

To determine if *Mll1* and *Mll2* deficiency during the retinal development displayed cell-type specific effects, retinal cross-sections of 1MO mutants and controls were immunostained with cell-type specific markers. Given ERG deficits in the *double CKO* mutants, we firstly assessed the staining of rod and cone Opsins (Figure 4A-C). Rhodopsin (Rho) staining was correctly localized to the OS in the *double KO* retina (Figure 4A). However, consistent with the results of H&E staining (Figure 3D), the length of Rho+ OS appeared shorter in the mutant than the *CreNeg* control (Figure 4A). Next, we stained for cone opsins. In mice, M and S cone subtypes were organized in dorsal-ventral counter-gradients, with M-opsin+ cones predominantly found in the dorsal retina and S-opsin+ cones in the ventral retina (Applebury et al., 2000). In the *double CKO* retinae, both M-opsin and S-opsin labeled cells were detected with the expected dorsal/ventral orientation (Figure 4B-C). However, while the number of M-opsin+ cells was reduced to about 20% of the control, no significant reduction of S-opsin+ cells was found (Figure 4H).

**Figure 4.**
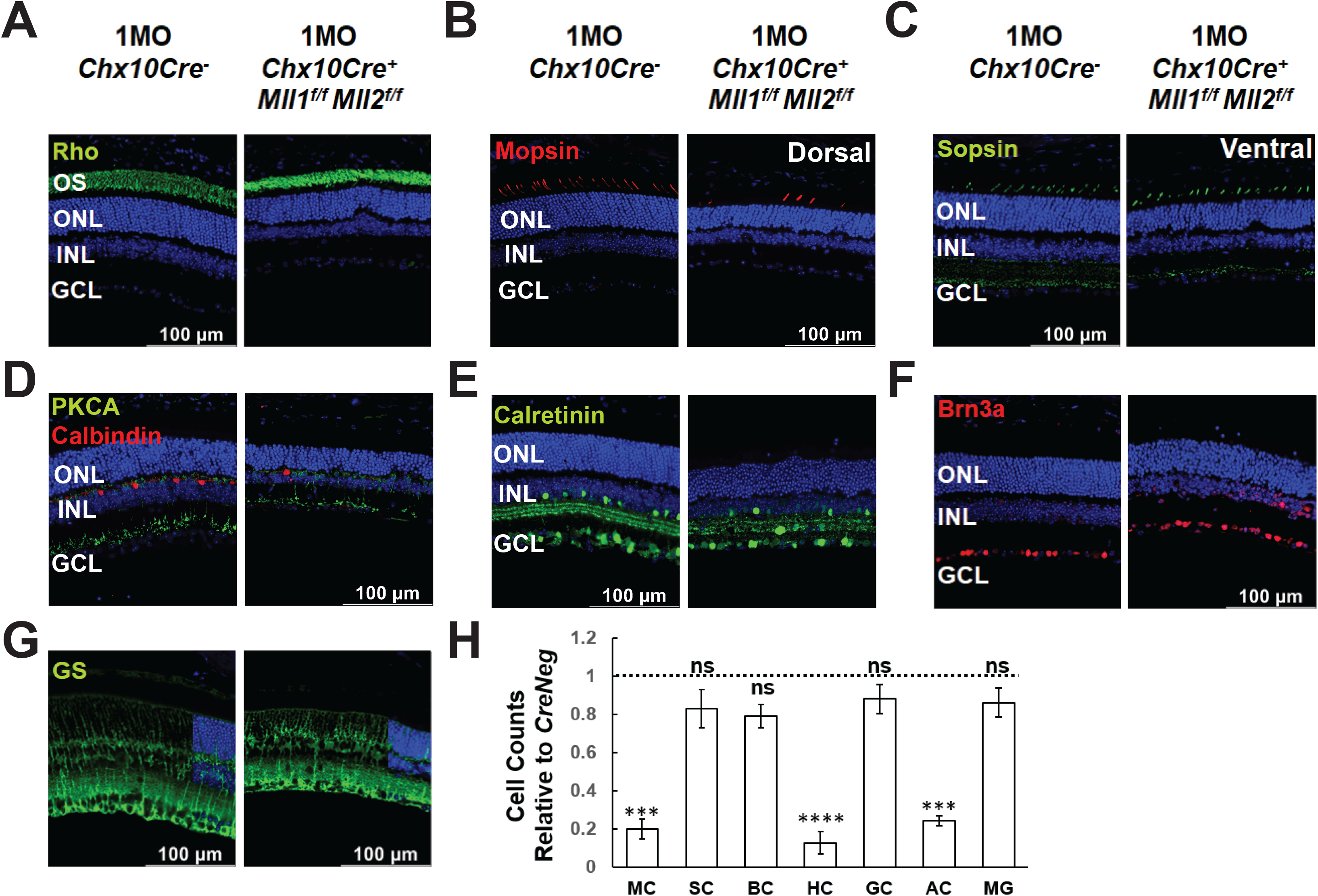
Altered cell type composition in *Chx10Cre*^+^*Mll1^f/f^Mll2^f/f^* at 1MO. (**A-G**) Immunostaining of the following cellular markers in 1MO retinae of the indicated genotypes: Rhodopsin (Rho, green in **A**) for rods, Medium-wave-sensitive opsin 1 (Mopsin, red in **B**) for M-cones, Short-wave-sensitive opsin 1 (Sopsin, green in **C**) for S-cones, Protein kinase C alpha (PKCA, green in **D**) for rod ON-bipolar cells, Calbindin D-28 (red in **D**) for horizontal cells, Brn3a (red in **E**) for ganglion cells, Calretinin (green in **F**) for amacrine and ganglion cells, and Glutamine synthetase (GS, green in **G**) for Müller glia. Nuclei are counterstained by DAPI (blue). Scale bar = 100µm for all image panels. (**H**) Cell counts for different retinal cell types in 1MO *Chx10Cre*^+^*Mll1^f/f^Mll2^f/f^* retinae, normalized to counts from *Chx10Cre^-^* (*CreNeg*) littermates. Error bars represent SEM (n≥3). Asterisks (***, ****) denote *p* ≤ 0.001, *p* ≤ 0.0001 respectively, ns means not significant by T-test. MC=M cone photoreceptors, SC=S cone photoreceptors, BC=bipolar cells, HC=horizontal cells, GC=retinal ganglion cells, AC=amacrine cells, MG=Müller glia.

Next, we assessed changes of neuronal cell types in the inner retina. The *double CKO* retinae displayed significantly decreased numbers (10-30% remaining) of Calbindin+ horizontal cells, and Calretinin+ amacrine cells (Figure 4D, E, H). In contrast, the numbers of PKCα+ rod-on bipolar cells (Figure 4D) and Brn3a+ ganglion cells (Figure 4F) in mutants generally remained comparable to the controls (Figure 4H). Interestingly, 1MO *single CKO* retinae also had decreased numbers of M-cones, horizontal cells, and amacrine cells (Supplemental Figure 4), but the degree of reduction was less severe than that in the *double CKO* retinae. Taken together, the distinct and redundant roles of MLL1 and MLL2 were essential for the integrity of M-cones, horizontal cells, and amacrine cells.

We also examined the number and morphological features of Müller glia by glutamate synthetase (GS) staining. GS+ Müller glia in the 1MO *double KO* retinae showed the typical cellular structure as compared to the control (Figure 4G), with cell bodies locating at the INL and endfeet terminating at the basal lamina. Mutant retinae also had similar numbers of Müller glia (Figure 4H), suggesting the development of these glial cells was generally unaffected in the mutants.

To determine if the loss of selected retinal cell types was due to defective cell type specification or maintenance, we examined the early cell types in P0 retinae by immunostaining (Supplemental Figure 5). We used antibodies targeting Retinoid X Receptor Gamma (RXRg) to label early fated cones, Onecut1 for early horizontal cells, Activating enhancer binding protein 2 alpha (AP2α) for early amacrine cells, and Paired box 6 (Pax6) for late-born RPCs (Supplemental Figure 5A). Cell count analysis showed that the numbers of these four cell types were comparable in the *double CKO* and *CreNeg* retinae (Supplemental Figure 5B), indicating that the production of early-born neurons and Pax6+ late RPCs was not severely affected in the *double CKO* retinae at P0.

We next analyzed the cell numbers of cones, horizontal and amacrine cells in mutant retinae at P14 when the cell fate specification of all retinal neurons was completed (Figure 5A). The cell numbers of these cell types in mutants were lower than *CreNeg* samples (Figure 5B), but the degree of reduction at P14 (20-50%) was less than that at 1MO (80-90%), indicating a progressive loss of cones, horizontal and amacrine cells in the *double CKO* retinae during the postnatal retinal development.

**Figure 5.**
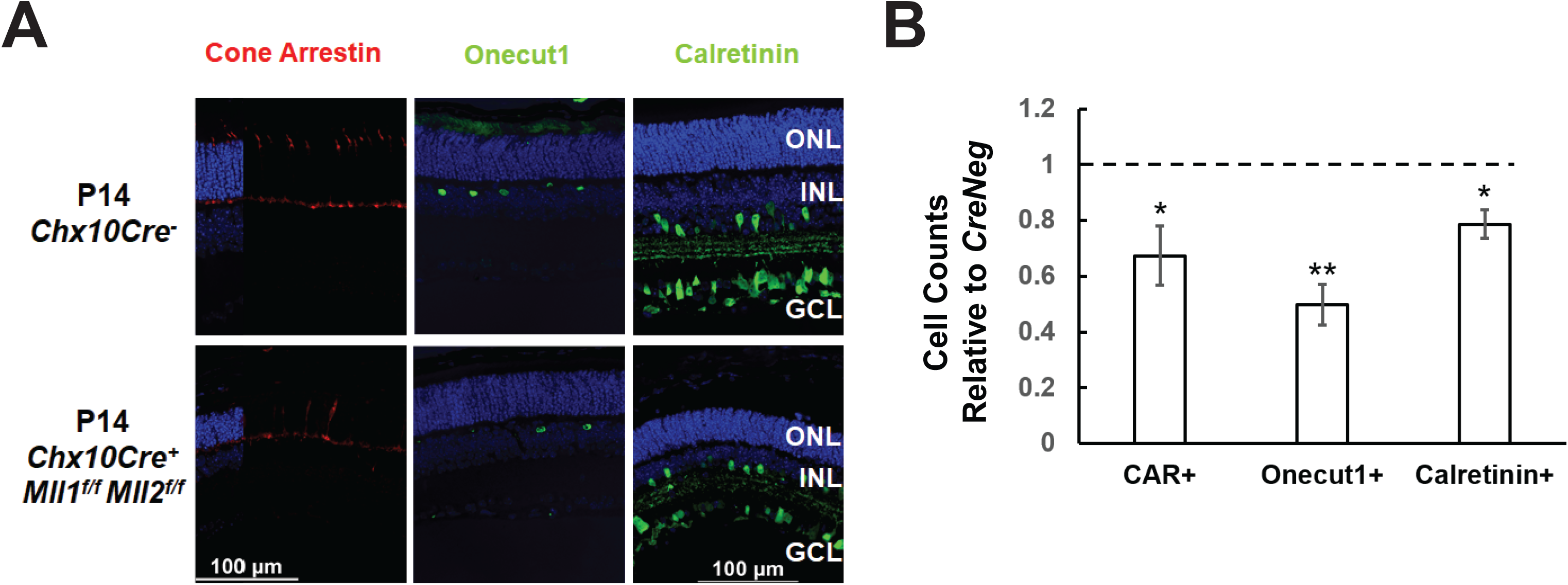
Altered cell type composition in *Chx10Cre*^+^*Mll1^f/f^Mll2^f/f^* at P14. (**A**) Immunostaining of cellular markers in P14 retinae: Cone arrestin (CAR, left panel, in red) for cones, Onecut1 (center panel, in green) for horizontal cells, and Calretinin (right panel, in green) for amacrine and retinal ganglion cells. Nuclei are counterstained by DAPI (blue). Scale bar = 100µm for all image panels. (**B**) Cell counts for different retinal cell types in P14 *Chx10Cre*^+^*Mll1^f/f^Mll2^f/f^* retinae, normalized to counts from *Chx10Cre^-^* (*CreNeg*) littermates. Error bars represent SEM (n≥3). Asterisks (*, **) denote *p* ≤ 0.05, *p* ≤ 0.01 respectively, ns means not significant by T-test.

To determine if programmed cell death contributed to the loss of neurons, we immunostained mutant retinae with activated caspase 3. At P0, a significant increase in the number of apoptotic cells (activated caspase 3+) were detected in mutant compared to *CreNeg* samples (Supplemental Figure 6A-B). Increased activated caspase 3+ cells were also notable at 1MO mutant retinae (Supplemental Figure 6C). We also assessed the expression of pro- and anti-apoptotic genes. The transcript levels of the pro-apoptotic gene *Bcl-2-associated X* (*Bax*) were upregulated in P14 *double CKO* retinae and more significantly in 1MO *double CKO* retinae relative to the *CreNeg* control, while the expression of the anti-apoptotic gene *B-cell lymphoma 2* (*Bcl2*) was insignificantly reduced in mutants (Supplemental Figure 6D). These data suggested that in *double CKO* retinae, the intrinsic apoptosis was initiated during the retinal development and accelerated at 1MO. This was consistent with retinal thinning and cell loss detected at P14 and 1MO, explaining the rapid degeneration afterwards.

To determine if *Mll1/Mll2* deficiency induced reactive gliosis, we performed immunostaining of Glial fibrillary acidic protein (GFAP) on retinal cross-sections of the *double CKO* mice. GFAP was the intermediate filament protein in retinal glia, its immunoreactivity indexed potential gliosis and correlation of neural damage. At 1MO, GFAP immunoreactivity was minimal in the control retinae, but was drastically intensified in the *double CKO* retinae (Figure 6A), indicating severe reactive gliosis. The GFAP+ reactive gliosis was lower in the *single CKO* retinae than that in the *double CKO* retinae. This high GFAP immunoreactivity in 1MO *double CKO* retinae was confirmed by GFAP qRT-PCR (data not shown). The GFAP immunoreactivity was comparable in the P14 *double CKO* retinae (Figure 6B) to the control. Thus, the glial activation in mutants likely occurred after eye-opening, accompanied by the loss of retinal integrity at both structural and functional levels.

**Figure 6.**
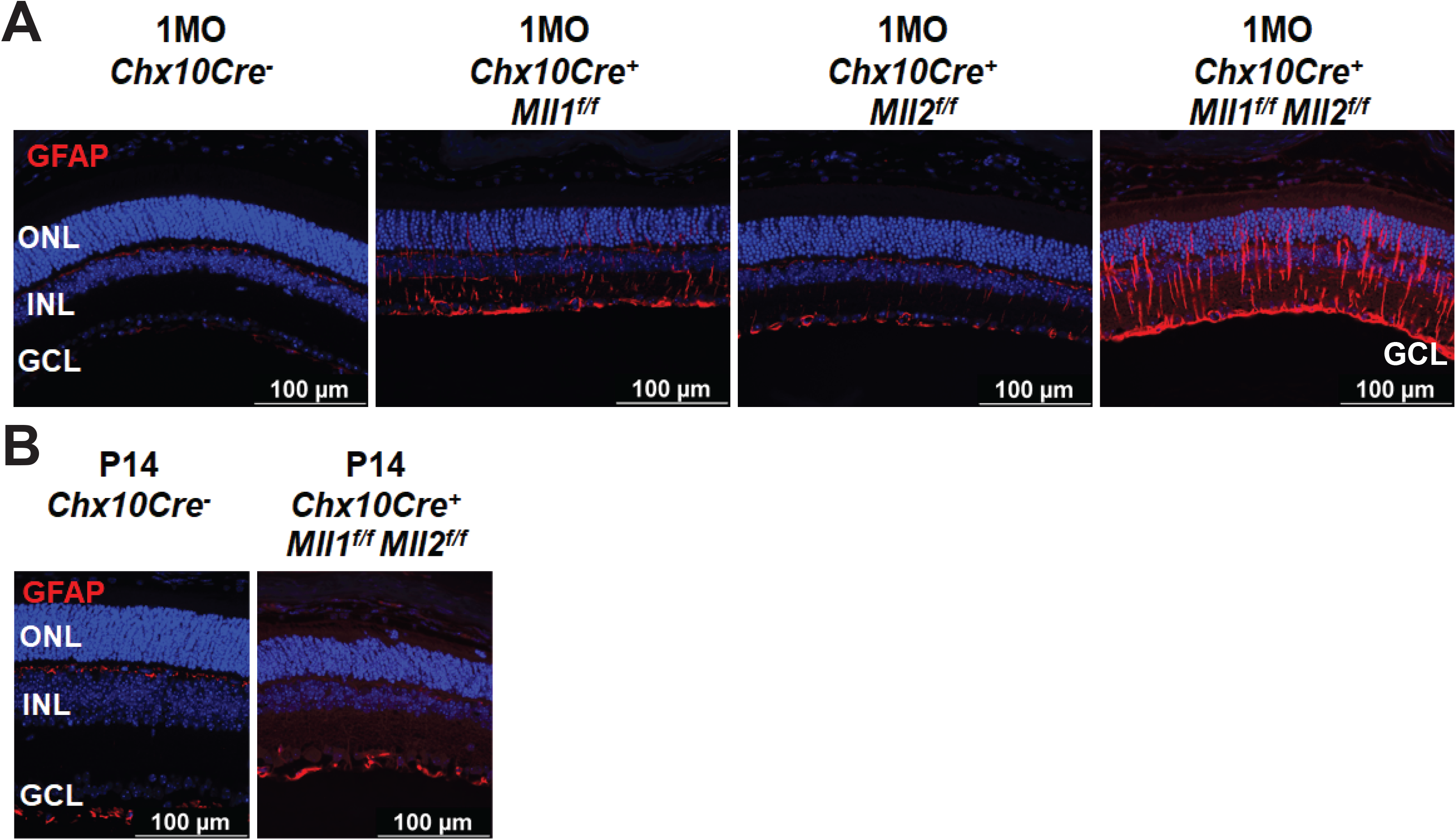
Glia activation in *Chx10Cre*^+^*Mll1^f/f^Mll2^f/f^*. (**A**) Glial fibrillary acidic protein (GFAP) immunostaining (in red) in 1MO mutant and control retinae. Nuclei are counterstained by DAPI (blue). Scale bar = 100µm for all image panels. (**B**) GFAP immunostaining in P14 mutant and control retinae.

### MLL1 and MLL2 regulate late progenitor cell proliferation in postnatal developing retina

Since cell proliferation defects were previously observed within P3 retinae of the *Chx10Cre*^+^*Mll1^f/f^* mutants (Brightman et al., 2018), we sought to determine if a similar phenotype could be found in the *double CKO* retinae at an early postnatal age. We performed Ki67 immunostaining for proliferating cells in the retinal cross-sections of P0 samples (Figure 7A). The number of Ki67+ proliferative cells in the *double CKO* retinae was decreased to 79% of control (Figure7B). We next examined if the cell cycle of Ki67+ proliferative cells was affected. Phospho-histone H3 (PH3) was used to label M-phase cells. PH3+ cells were found at the outer retinal neuroblast layer (NBL) of the *double CKO* and *CreNeg* retinae (Figure 7A). We did not find a significant difference in Ki67+ and PH3+ co-labelled cells between these samples. To detect S-phase cells in P0 retinae, 5-Ethynyl-2′-deoxyuridine (EdU) was IP-injected into P0 mouse pups, and retinae were harvested 4 hours later (Figure 7A right panel). The *double CKO* retinae possessed fewer EdU+ cells than *CreNeg* control (Supplemental Figure 7A, 7C). More importantly, the *double CKO* retinae had a decrease in Ki67+/EdU+ co-labelled cells, down to 72% of the control (Figure 7B). These data suggested that *Mll1* and *Mll2* deficiency affected cell proliferation and cell cycle dynamics in P0 retinae.

**Figure 7.**
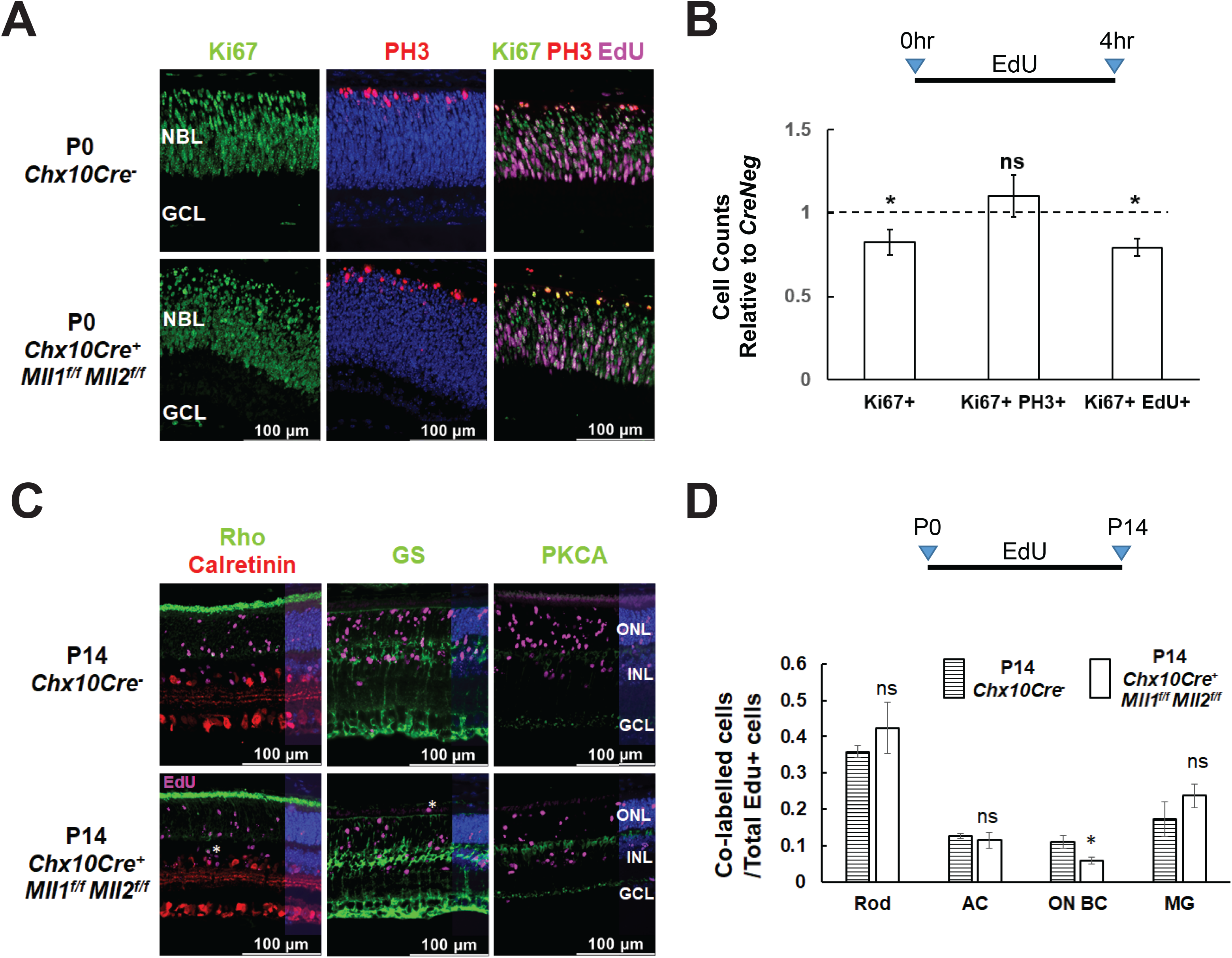
Cell proliferation defects in *Chx10Cre*^+^*Mll1^f/f^Mll2^f/f^*. (**A**) Immunostaining of cell proliferation and cell cycle markers in the indicated P0 retinae: Ki67 (left panel, in green) for proliferating cells, Ki67/PH3 (center panel, Ki67 in green, PH3 in red) for cells in the M phase, Ki67/EdU (right panel, Ki67 in green, EdU in magenta) for cells in the S phase. Nuclei (center panel) are DAPI counterstained in blue. Scale bar = 100µm for all image panels. (**B**) Cell counts for proliferating cells in P0 *Chx10Cre*^+^*Mll1^f/f^Mll2^f/f^* retinae, normalized to counts from *Chx10Cre^-^* (*CreNeg*) littermates. Error bars represent SEM (n=3). Ki67+=Ki67 labelled cells, Ki67+PH3+=Ki67 and PH3 co-labelled cells, Ki67+EdU+=Ki67 and EdU co-labelled cells. (**C**) P14 retinal immunostaining of Rho/Calretinin/EdU (left panel, Rho in green, Calretinin in red, EdU in magenta), GS/EdU (center panel, GS in green, EdU in magenta), PKCA/EdU (right panel, PKCA in green, EdU in magenta). White asterisks indicate mislocalized cells. (**D**) Cell compositions of co-labelled cells in P14 retinae of the indicated genotypes. Error bars represent SEM (n=3). Results of statistical analysis are shown. Asterisks (*) denote *p* ≤ 0.05, ns means not significant by T-test. Rod=rod photoreceptors, AC=amacrine cells, ON BC=ON bipolar cells, MG= Müller glia.

To determine impacts of *Mll1*&*Mll2* deficiency on the production of specific cell types from late-born retinal progenitor cells (RPC), we performed EdU pulse-chase experiments. EdU-labeled P0 proliferating cells were assessed for specified cell-types at P14 (Figure 7C, D). Firstly, at P14, there were fewer EdU+ cells in the *double CKO* retinae than in *CreNeg* controls (Supplemental Figure 7B, C). Secondly, the percentage of EdU co-labelled ON-bipolar cells in overall EdU-labelled cells was reduced about one half as compared to the control, but those of rods, late-born amacrine cells and Müller glia remained comparable (Figure 7C, D). These results suggested that, despite moderately reduced ON-bipolar cell production from P0 progenitor cells, overall, the intrinsic neurogenesis program was not severely compromised in the *double CKO* retinae.

### *Mll1* and *Mll2* deficiency reduces the expression of phototransduction genes without altering the promoter occupancy of the active histone mark H3K4me3

To unveil the molecular mechanisms for the cellular and functional defects in the *double CKO* retinae, we profiled specific gene expression changes at P14 and 1MO. qRT-PCR analysis showed there was a general trend toward reduced expression with tested transcripts in *double CKO* retinae as compared to the *CreNeg* retinae, suggesting that *Mll1*/*Mll2* deficiency had a negative impact on gene expression in general (Figure 8A). However, the transcript level of *lysine demethylase 6B* (*Kdm6b*) was slightly upregulated in 1MO *double CKO* retinae. The downregulation of the cell-type specific transcript levels supported the results of immunohistochemistry (Figures 4, 5), further confirming cell composition changes at these two ages. In particular, this downregulation was most prominent in three cell types (Figure 8A), M-cones (*Opn1mw*, 30-50%), horizontal cells (*Prox1*, ∼60%) and amacrine cells (Pax6, ∼60%). Lastly, the changes in transcript expression at 1MO were more profound than those at P14, which was consistent with the progressive loss of the affected neurons. For example, the relative expression of *Opn1mw* was reduced from 52% at P14 to 34% at 1MO. The relative expression of *Rho* decreased from insignificant 70% at P14 to significant 48% at 1MO. These results, combined with the reduction of rod OS length at 1MO, suggested that rod cell integrity was affected before degeneration peaks. Additionally, we performed qRT-PCR with early-born cell markers at P0, including *Thrb* and *Rxrg* (cones), *Onecut1* (horizontal cells), *Chx10* and *Pax6* (RPCs). No expression changes were detected in P0 *double CKO* retinae relative to the control (Supplemental Figure 8), suggesting that early RPCs and their neurogenesis were unaffected by *Mll1*/*Mll2* deficiency.

**Figure 8.**
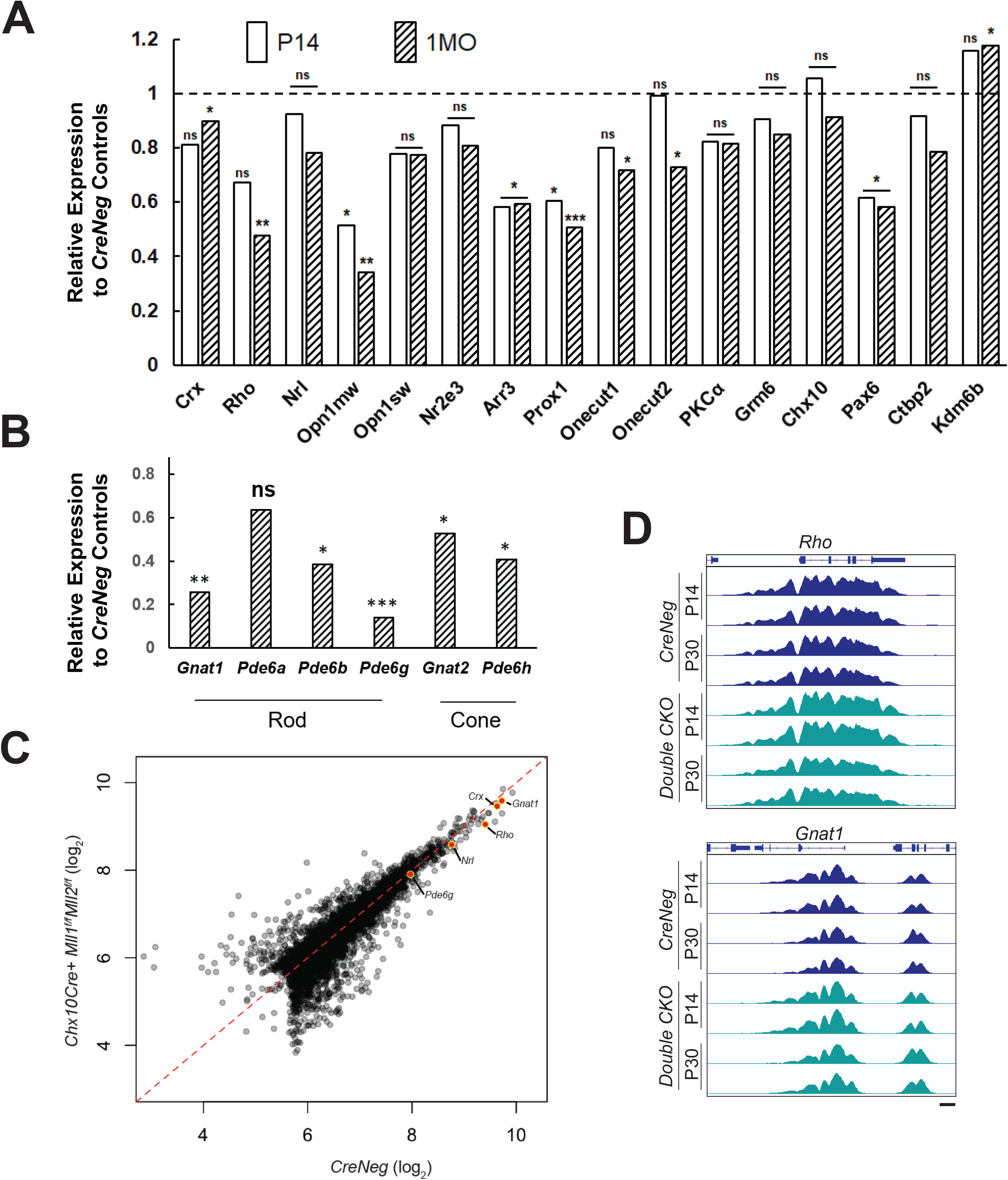
Gene expression profile in P14 and 1MO *Chx10Cre*^+^*Mll1^f/f^Mll2^f/f^*. (**A**) qRT-PCR analysis of selected genes in P14 and 1MO *Chx10Cre*^+^*Mll1^f/f^Mll2^f/f^* retinae. Results are plotted as relative expression to *Chx10Cre^-^* (*CreNeg*) littermate controls (n≥4). (**B**) qRT-PCR analysis of phototransduction gene expression in 1MO *Chx10Cre*^+^*Mll1^f/f^Mll2^f/f^* retinae. Results are plotted as relative expression to *Chx10Cre^-^* (*CreNeg*) littermate controls (n≥4). Asterisks (*, **, ***) denote *p* ≤ 0.05, *p* ≤ 0.01, *p* ≤ 0.001 respectively by one-way ANOVA with Tukey’s multiple comparisons, and ns means not significant. (**C**) Quantitative comparison of data from all H3K4me3 peaks in 1MO *CreNeg* and *Chx10Cre*^+^*Mll1^f/f^Mll2^f/f^* datasets. Promoter peaks of key phototransduction genes are highlighted. (**D**) ChIP-Seq data from P14 and 1MO *Chx10Cre*^+^*Mll1^f/f^Mll2^f/f^* and *CreNeg* retinae for H3K4me3 shows no differences at key rod phototransduction loci.

We also examined gene expression of several other components in the rod and cone phototransduction cascades. qRT-PCR was performed to analyze transcripts of *G protein subunit alpha transducin 1* (*Gnat1*), *transducin 2* (*Gnat2*), *Phosphodiesterase 6a* (*Pde6a*), *6b* (*Pde6b*), *6g* (*Pde6g*), *6h* (*Pde6h*). All tested genes besides *Pde6a* showed significant reduction in 1MO *double CKO* retinae (Figure 8B). Notably, the relative expression of *Gnat1* and *Pde6g* was decreased to 25% and 14% of the control, respectively. These results suggested that the rod and cone phototransduction cascade was disrupted in 1MO *double KO* retinae, providing an explanation for the lack of ERG responses. In contrast, the *single CKO* samples only had moderate or insignificant changes in gene expression (Supplemental Figure 9).

Since MLL1 and MLL2 function to methylate H3K4 at active gene promoters, we hypothesized that the gene misregulation in the mutants may be caused by the lack of the active histone mark H3K4me3 at photoreceptor specific promoters. We performed whole-retina chromatin immunoprecipitation sequencing (ChIP-Seq) to compare the *double CKO* mutants with *CreNeg* controls. Because rods represented 70% of all retinal cells, ChIP data represented a primarily rod signal in bulk ChIP-Seq analysis of P14 and 1MO samples (before severe rod degeneration in the mutants). Biological replicates displayed >98% concordance, indicating excellent reproducibility (Supplemental Figure 10A). Next, we compared the ChIP signal between mutants and controls and identified differentially enriched peaks at each age as well as those peaks that change over time in each model (Supplemental Figure 10B). To our surprise, there were only 279 peaks that displayed significant differences (>2 fold-change, P< 0.05) in these comparisons (Figure 8C). To understand the normal deposition of these histone modifications in retinal development, we analyzed the average change in signal over retinal development (E14-P21) (Aldiri et al., 2017). Peaks that were decreased in mutants generally showed consistent H3K4me3 across development, while peaks gained in mutants showed very little deposition of any active marks in the controls. We inspected the profiles of H3K4me3 at the rod genes that showed dramatic gene expression reductions (Figure 8B) but did not observe concordant reduction in the presence of H3K4me3 (Figure 8D). Together, these results suggested that *Mll1/Mll2* deficiency did not affect the remodeling of photoreceptor chromatin and the deposition of the H3K4me3 mark on promoters activated during cell type specification; other mechanisms must be responsible for the reduced expression of photoreceptor-specific genes.

### MLL1 and MLL2 are required for the maintenance of fated M-cones, but not rods

Although defective rod and cone phenotypes were seen in *Chx10-Cre*-mediated *double CKO* retinae, it was unclear if these phenotypes were stemmed from primary and/or secondary effects of *Mll1/Mll2* deficiency. To address this question, we determined cell autonomy by investigating the effects of *Mll1*/*Mll2* deficiency in fated rods and cones, respectively. To ablate *Mll1/Mll2* functions in fated rods, we crossed *Mll1^f/f^Mll2^f/f^* mice to *Nrl-Cre* mice (Brightman et al., 2016), which began to express Cre recombinase in developing rods soon after rod cell specification and sustained Cre expression in adult rods. We also ablated *Mll1/Mll2* in fated M-cones by crossing *Mll1^f/f^Mll2^f/f^* mice to *Mcone-Cre* (*HRGP-Cre*) mice (Le et al., 2004), which expressed Cre recombinase in M-opsin+ cones beginning at early postnatal ages. H&E staining of 1MO *NrlCre*^+^*Mll1^f/f^Mll2^f/f^* and *MconeCre*^+^*Mll1^f/f^Mll2^f/f^* showed no apparent morphological abnormality (Figure 9A).

**Figure 9.**
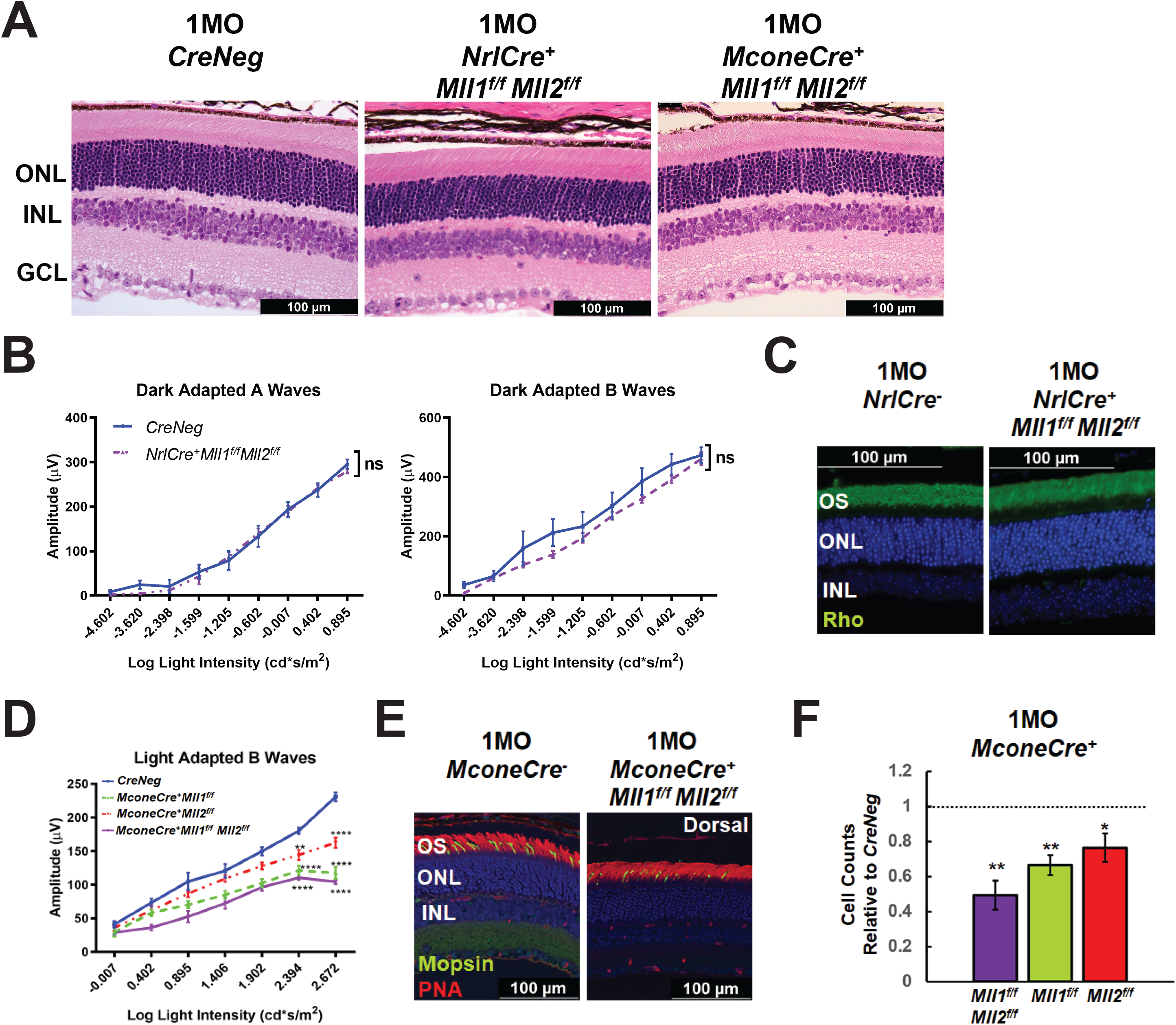
Functional and morphological assessments of 1MO *NrlCre*^+^*Mll1^f/f^Mll2^f/f^* and *MconeCre*^+^*Mll1^f/f^Mll2^f/f^*. (**A**) H&E cross-section staining of 1MO *Cre^-^* (*CreNeg,* left panel), *NrlCre*^+^*Mll1^f/f^Mll2^f/f^* (center panel) *MconeCre*^+^*Mll1^f/f^Mll2^f/f^* (right panel) retinae. Scale bar = 100µm for all image panels. (**B**) Dark adapted A-wave (left) and Dark adapted B-wave (right) ERG analysis of 1MO *NrlCre*^+^*Mll1^f/f^Mll2^f/f^* and *Cre^-^* (*CreNeg*) mice. Mean amplitudes (µV) are plotted against stimulus light intensity. Error bars represent SEM (n ≥ 4). (**C**) Rho immunostaining in 1MO *NrlCre^-^* and *NrlCre*^+^*Mll1^f/f^Mll2^f/f^* retinae. (**D**) Light adapted B-wave ERG analysis of 1MO *MconeCre*^+^*Mll1^f/f^Mll2^f/f^*, *MconeCre*^+^*Mll1^f/f^*, *MconeCre*^+^*Mll2^f/f^*, and *Cre^-^* (*CreNeg*) mice. Mean amplitudes (µV) are plotted against stimulus light intensity. Error bars represent SEM (n ≥ 4). (**E**) Mopsin and PNA immunohistochemical staining in 1MO *MconeCre^-^* and *MconeCre*^+^*Mll1^f/f^Mll2^f/f^* dorsal retinae. (**F**) Cell counts for M cone photoreceptors in 1MO *MconeCre*^+^*Mll1^f/f^Mll2^f/f^ MconeCre*^+^*Mll1^f/f^*, and *MconeCre*^+^*Mll2^f/f^* retinae, normalized to counts from *MconeCre^-^* (*CreNeg*) littermates. Error bars represent SEM (n=3). All statistics is done by two-way ANOVA and Tukey’s multiple comparisons. Asterisks (*, **, ***) denote *p* ≤ 0.05, *p* ≤ 0.01, *p* ≤ 0.001 respectively, ns means not significant.

We next measured ERG responses in 1MO *NrlCre*^+^*Mll1^f/f^Mll2^f/f^* mice. No defects in dark-adapted A and B waves were detected in mutants compared to the controls (Figure 9B), and light-adapted ERG was also comparable (data not shown). Furthermore, Rho immunostaining showed no significant difference in OS morphology and thickness (Figure 9C). Transcript-level expression analysis of rod-enriched genes by qRT-PCR detected no changes in 1MO mutant retinae relative to the control (Supplemental Figure 11A). We did not detect any morphological and functional changes up to 6MO (data not shown). Collectively, these results suggested that *Mll1*/*Mll2* deficiency in developing and mature rods has limited impacts on cellular structure, function and survival. Thus, MLL1 and MLL2 functions were not required for the maintenance of fated rods.

By comparison, *MconeCre*^+^*Mll1^f/f^Mll2^f/f^* mice displayed cone functional and morphological defects. The amplitudes of light-adapted B waves of 1MO *MconeCre*^+^*Mll1^f/f^Mll2^f/f^* mice were significantly decreased as compared to controls (Figure 9D). 1MO *MconeCre^+^Mll1^f/f^* and *MconeCre^+^Mll2^f/f^* mice also showed decreased ERG responses as compared to controls, but were less severe than *MconeCre*^+^*Mll1^f/f^Mll2^f/f^* mice (Figure 9D). These results suggested that MLL1 and/or MLL2 were required for the functional maintenance of M-cones even after the cone differentiation.

We next performed immunostaining of Opn1mw (M-cones) together with PNA (all cones), or cone arrestin (Arr3) in retinal cross-sections of 1MO *MconeCre^+^Mll1^f/f^*, *MconeCre^+^Mll2^f/f^*, *MconeCre*^+^*Mll1^f/f^Mll2^f/f^* retinae (Figure 9E-F, Supplemental Figure 12). Quantification showed that the number of Opn1mw+ cones in *MconeCre*^+^*Mll1^f/f^Mll2^f/f^* retinae was reduced to 50% of the control, while the numbers in *MconeCre^+^Mll1^f/f^* or *MconeCre^+^Mll2^f/f^* was down to only 70-80% (Figure 9F). To determine any spatial patterns associated with the cone number reduction, we performed M-opsin and PNA co-staining on 1MO whole-mount *MconeCre*^+^*Mll1^f/f^Mll2^f/f^* retinae (Supplemental Figure 13A). As expected, in 1MO control samples, Opn1mw+ cones were largely localized to the dorsal retina (Supplemental Figure 13A). By comparison, fewer Opn1mw+ cones were found in the dorsal region of *MconeCre*^+^*Mll1^f/f^Mll2^f/f^* retinae (Supplemental Figure 13A). Cell count analysis showed that nearly half of Opn1mw+ (PNA+) cells were absent in the dorsal mutant retinae, while the cone cell numbers were comparable in the ventral retinae (Supplemental Figure 13B), confirming *Mll1*/*Mll2* deficiency in cones specifically affected M-cone integrity. qRT-PCR analysis showed significantly decreased expression of *Opn1mw* and *Arr3* in 1MO *MconeCre*^+^*Mll1^f/f^Mll2^f/f^* retinae, but not in *MconeCre^+^Mll1^f/f^* or *MconeCre^+^Mll2^f/f^* retinae (Supplemental Figure 11B-C), further confirming the immunostaining results. Taken together, these results described the cell autonomous effect of *Mll1*/*Mll2* deficiency on M-cone functional maintenance.

## DISCUSSION

### Redundant roles of MLL1 and MLL2 in retinal development and maintenance

MLL1 and MLL2 have been reported to play non-overlapping roles in development and cancer (Chen et al., 2017). In this manuscript, we used the conditional knockout strategy in mice to provide several pieces of evidence supporting that MLL1 and MLL2 redundant functions were required for the development and maintenance of neuronal function in the retina.

When MLL1 or MLL2 was removed from Chx10+ retinal progenitor cells, similar phenotypic defects were observed, including moderately reduced photoreceptor responses to light stimuli (Figure 2, Supplemental Figure 1), loss of M-cones, horizontal and amacrine cells (Supplemental Figure 4). Thus, MLL1 and MLL2 essentially functioned in the indistinguishable regulatory pathways during retinal development and maintenance. MLL1 contributed more than MLL2 in each pathway, as all observed phenotypes were more severe in *Chx10Cre*^+^*Mll1^f/f^* than *Chx10Cre*^+^*Mll2^f/f^*. In each *single CKO* retinae, there were no decreases or compensatory increases of transcript levels of the other *Mll* gene.

When both MLL1 and MLL2 were removed, all phenotypic defects found in the *single CKO* retinae became much more severe. Notably, the *double CKO* mutants produced null ERG responses along with severely affected phototransduction gene expression (Figures 2, 8), massive reactive gliosis (Figure 6) and apoptosis (Supplemental Figure 6) at 1MO and rapid retinal degeneration after 1MO (Figure 3). However, compound heterozygotes of *Mll1* and *Mll2* mutants produced significantly less severe ERG defects than the *double CKO* mutants, which was comparable to those in *single CKO* mutants (Supplemental Figure 1B-C). These results collectively confirmed that MLL1 contributed more than MLL2 in their redundant functions.

The functional redundancy between MLL1 and MLL2 was specific to this pair of MLLs. We previously reported that the *Mll1/Mll3 double CKO* retinae produced a phenotype similar to that seen in *single Mll1 CKO* retinae (Brightman et al., 2018). MLL1 or MLL2 formed a complex distinct from those formed by other MLL family members, which provided a possible explanation for the redundant roles of MLL1 and MLL2. We proposed MLL1 and MLL2 acted together to regulate the neuronal proliferation, development and survival in the retina (Figure 10). Although we emphasized the redundant functions of MLL1 and MLL2 in this study, we did not rule out distinct functions of each gene, particularly for unique mechanisms of action at the molecular level. As an example, previous studies reported both MLL1 and MLL2 regulated *Hox* gene expression essential for body patterning during development, but they bound on distinct *Hox* loci with MLL1 on *HoxA* and *HoxC* and MLL2 on *HoxB* (Glaser et al., 2006; Milne et al., 2002; Yu et al., 1995). Future studies are needed to determine the molecular mechanisms underlining MLL1/MLL2 redundant functions in the retina, including their individual actions on specific neurons.

**Figure 10.**
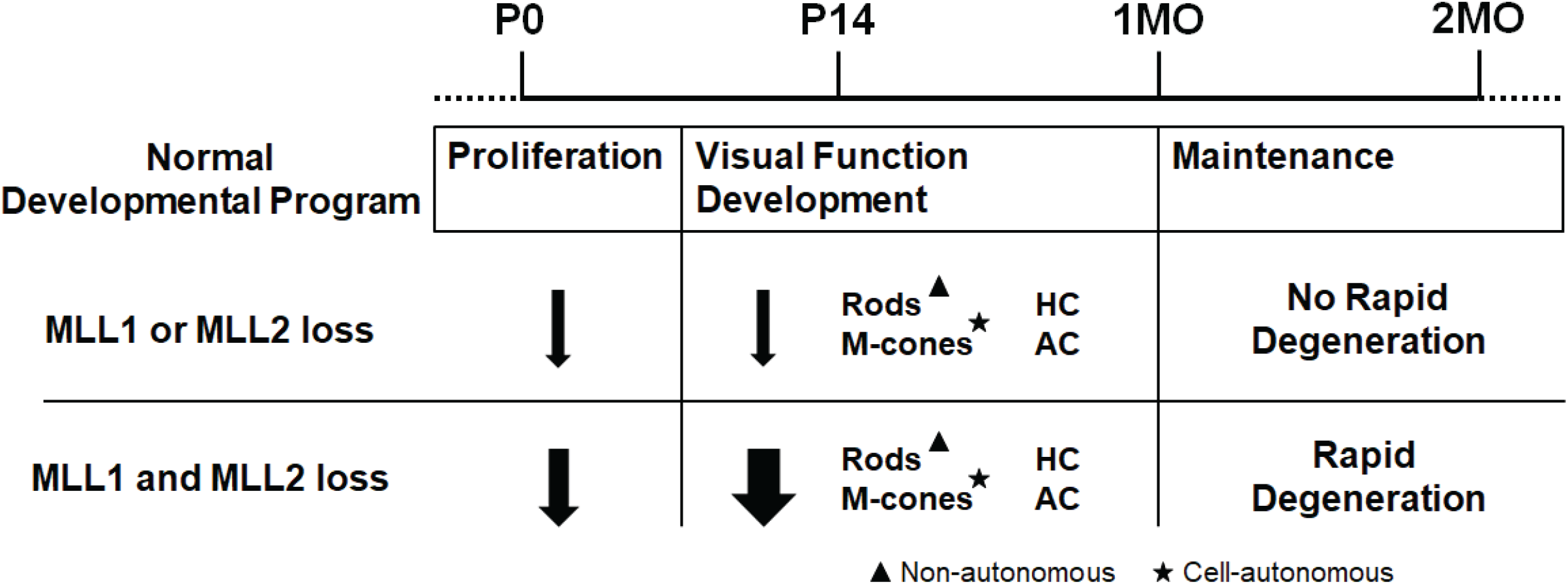
Summary of retinal defects in *Mll1* and *Mll2* conditional knockouts.

### The mechanisms underlying retina layer thinning and functional deficits in *Mll1/Mll2* double *CKO* samples

One major phenotype of *Mll1/Mll2* deficiency was the thinning of retinal layers and the loss of specific types of neurons. In P0 *Chx10Cre*^+^*Mll1^f/f^Mll2^f/f^* retinae, the numbers of early-born neurons and expression levels of specific transcription factors were comparable to controls. Thus, the genesis and cell specification of the early-born neurons were normal. However, the development and maintenance of rods, M-cones, horizontal and amacrine cells were negatively affected at P14 (Figure 5) and 1MO (Figure 4), along with increased apoptosis throughout the first 4 weeks of postnatal development (Supplemental Figure 6).

Both cell autonomous and non-autonomous mechanisms were involved. The cellular mechanisms for the loss of rods were more complex, and likely involve cell non-autonomous (secondary) effects. Key observations included, firstly, the majority of rods survived between P14 to 1MO with shorter segments. At 1MO, apoptosis was apparent to cells at INL, not to ONL cells. Secondly, significant injury-induced reactive gliosis was present in the 1MO *Chx10Cre*^+^*Mll1^f/f^Mll2^f/f^* retinae, correlating cellular stress in general. Lastly and most importantly, the removal of both *Mll1/Mll2* for developing rods (by *Nrl-Cre*) did not produce rod functional deficits or cell loss. In addition, rod degeneration as secondary effects by neuronal loss at INL has been reported. For example, the loss of horizontal cells due to Onecut1 and Onecut2 TF deficiency induced subsequent rod degeneration (Sapkota et al., 2014).

In contrast, the M-cone phenotype was largely attributed to a cell autonomous mechanism, as M-cone defects was observed in retinae with *Chx10-Cre* and *Mcone-Cre* mediated *Mll1/Mll2* deficiency. Notably, *Mll1/Mll2* deficiency only affected the M-cones, not S-cones. The mechanisms for such specificity of the MLL actions on cone subtypes remain to be determined. Further studies are needed to determine if protein factors regulating the S/M cone gradient or cone opsin expression are affected in the *double CKO* mutants.

Defects in neurogenesis may also contribute to retinal thinning and reduced cell numbers in INL of the *double CKO*, particularly during the first two weeks of postnatal development. P0 *Chx10Cre*^+^*Mll1^f/f^Mll2^f/f^* retinae had reduced number of Ki67+ and EdU+ co-labelled cells, suggesting *Mll1* and *Mll2* collectively affect S-phase entry of retinal proliferating cells. This was consistent with published studies from mouse brain tissue and human cancer cell lines (Chen et al., 2017; Denissov et al., 2014; Jakovcevski et al., 2015; Kerimoglu et al., 2017). In our previous study, *Chx10Cre*^+^*Mll1^f/f^* showed an insignificant decrease in S-phase cell number, while this study presented a further decline in the *double CKO* retinae, suggesting a potential redundancy of *Mll1* and *Mll2* in regulating S-phase entry of retinal progenitors. Our EdU pulse-chase experiments at P0 to P14 showed that after cell cycle exit, P0 progenitor cells in the *double CKO* retinae could produce all four types of late-born cells, namely, rods, ON-bipolar cells, amacrine cells and Muller glial. Together with the unchanged number of early-born neurons at P0, these results suggested that *Mll1* and *Mll2* did not regulate retinal cell-type specification. Although there was a reduction of ON-bipolar cells in EdU pulse-chase experiments from P0 to P14, the overall bipolar cell numbers were unchanged at 1MO. Further studies are required for examining the functions of MLLs in different windows of the retinal development.

1MO *Chx10Cre*^+^*Mll1^f/f^Mll2^f/f^* mutants displayed null ERG response, despite the presence of rows of ONL cells. One possible explanation was the loss of synaptic connections between photoreceptors and inner neurons. The presynaptic proteins Ctbp2/RIBEYE and VGLUT1 were present at the outer plexiform layer (Supplemental Figure 14) of the *double CKO* retinae, although we could not conclude if synaptic connections were functional. The most likely explanation was the decrease in phototransduction, based on two observations: 1) the absence of A-waves indicating no signals of ERG response, and 2) the downregulation of key rod and cone phototransduction genes. It was unclear if other mechanisms affecting phototransduction were involved.

It remains to be determined if gene misregulation in specific neurons of *Mll1/Mll2* deficient retinae is due to altered histone H3 K4-methylation. We attempted to address this question by examining changes of rod-specific gene expression and chromatin occupancy of the H3K4me3 mark. Our surprising finding was that in spite of the downregulation of rod-specific genes, normal H3K4me3 signals were detected at the regulatory regions of these genes in *Chx10-Cre* mediated *Mll1/Mll2* knockouts. Instead, H3K4me3 changes were seen in the regulatory regions of constitutively expressed genes that were unknown to have rod-specific functions. Thus, both H3K4me3-dependent and independent mechanisms may attribute to the misregulation in rod functional development and maintenance. It is possible that the misregulation mechanisms involved in minor cell types, such as M-cones, horizontal and amacrine cells might be different from those of rods. Cell type-specific knockout and single cell-related approaches are needed to elucidate these mechanisms.

## CONCLUSIONS

In conclusion, this study is the first kind to address the redundancy functions of MLLs in neuroretinal development and maintenance. Our findings highlight the complex functions of the MLL1 and MLL2 in neuronal cell types. Our findings may inform the pathogenesis of the syndromic neurological disorders associated with human *MLL1* and *MLL2* mutations and phenotype variability.

## MATERIALS AND METHODS

### Transgenic mouse lines

All mice in this study were on the genetic background of *C57BL/6J* (the Jackson Laboratory, Stock No: 000664) free of *rd1* and *rd8* mutations. The *Cre*, *Mll1 flox*, *Mll2 flox* mouse lines were obtained from various published colonies. Both male and female mice were used in experiments.

All animal procedures were conducted according to the Guide for the Care and Use of Laboratory Animals of the National Institute of Health, and were approved by the Washington University in St. Louis Institutional Animal Care and Use Committee.

### Histology and immunohistochemistry (IHC)

Eyes were enucleated at various ages with the superior/dorsal side marked on the cornea, and fixed in 4% paraformaldehyde at 4°C overnight for paraffin embedded sections. Each retinal cross-section was cut 5 microns thick on a microtome. Hematoxylin and Eosin (H&E) staining was performed to examine retinal morphology. For IHC staining, sections firstly went through antigen retrieval with citrate buffer, and blocked with a blocking buffer of 5% donkey serum, 1% BSA, 0.1% Triton-x-100 in 1X PBS (pH-7.4) for 1 hour. Sections were then incubated with primary antibodies at 4°C overnight. Sections were washed with 1X PBS containing 0.01% TritonX-100 (PBST) for 30 minutes, and then incubated with specific secondary antibodies for 1 hour. Primary and secondary antibodies (Supplemental Table 1) were applied with optimal dilution ratios. All slides were mounted with hard set mounting medium with DAPI (Vectashield, Vector Laboratories, Inc., CA).

For IHC staining of retinal cryo-sections, eyes were fixed in 4% paraformaldehyde after the removal of lens for 1 hour. Eyes were then immersed in increasing concentrations of phosphate-buffered sucrose solutions, and incubated overnight at 4 °C in 20% phosphate-buffered sucrose solution. Eyes were incubated in a 1:1 mixture of OCT (optimal cutting temperature; Sakura Finetek) and 20% phosphate-buffered sucrose for 1 hour. Eyes were embedded in OCT and snap frozen. Blocks were sectioned at 5 μm. Blocking and antibody staining were processed as described above.

For IHC staining of whole flat-mount retinae, eyes were fixed in 4% paraformaldehyde for at least 6 hours. Retinae were dissected and blocked in the blocking buffer for 2 hours. Retinae were incubated with primary antibodies at 4°C overnight, followed by 1X PBST wash for 10 minutes, and then incubated with secondary antibodies for 3 hours. Upon a final 1X PBST wash for 10 minutes, retinae were placed flat on slides. All images were taken on a Leica DB5500 microscope.

### EdU labelling

Edu labelling was performed with the Click-iT EdU Assay Kit (Invitrogen, ThermoFisher Scientific). Mice were IP injected with 10μL of 10mM Edu per gram mouse weight at P0. Eyes were fixed and paraffin embedded with procedures described above. EdU staining was done according to the manufacturer’s protocol. IHC staining was done prior to EdU staining for experiments involved both steps.

### Morphometry and cell count analysis

Thickness of retinal layers were measured on the H&E images at specific locations from the optical nerve head. Results of measurements were plotted in a spider graph. For cell counting analysis, numbers of fluorescent objects were tallied in the entire cross-section. At least 4 biological replicates of each genotype were used in the statistical analysis. Two-way ANOVA with multiple comparisons or Student’s t-test were performed with P<0.05, CI:95% using Graphpad Prism 8 (GraphPad Software, CA).

### Electroretinogram (ERG)

Whole animal ERGs were performed on 1- and 2-month-old mice using UTAS-E3000 Visual Electrodiagnostic System with EM for Windows (LKC Technologies Inc., MD). Mice were dark-adapted overnight prior to the tests. The experimental procedures were adapted from the previously published study of our lab (Brightman et al., 2018). ERG responses of biological replicates were recorded, averaged and analyzed using Graphpad Prism 8 (GraphPad Software, CA). The mean peak amplitudes of dark-adapted A and B waves and light-adapted B waves were plotted against log values of light intensities (cd*s/m^2^). The statistical analysis was done by two-way ANOVA with multiple pairwise comparisons (Tukey’s).

### Quantitative PCR (qPCR)

Each RNA sample was extracted from 2 retinae of a mouse using the NucleoSpin RNA Plus kit (Macherey-Nagel, PA). RNA concentrations were measured using a NanoDrop One spectrophotometer (ThermoFisher Scientific). 2μg of RNA was used to produce cDNA using First Strand cDNA Synthesis kit (Roche, IN). Technical triplicates were run for each gene. Primers used in this study were listed in Supplemental Table 2. The reaction master mix consisted of EvaGreen polymerase (Bio-Rad Laboratories, CA), 1μM primer mix, and diluted cDNA samples. Samples were run using a two-step 40-cycle protocol on a Bio-Rad CFX96 Thermal Cycler (Bio-Rad Laboratories, CA). Data were analyzed with QBase software (Biogazelle, Belgium). The statistical analysis was done by two-way ANOVA or Student’s t-test with *p*<0.05, CI:95% using Graphpad Prism 8.

### Chip-Seq

Chromatin immunoprecipitation (ChIP) assay was performed using six pooled P14 / P30 *Mll1&2 KO*, or (*Chx10)* CreNeg control retinas per sample as previously described (Tran et al., 2014). Each sample was dissected and chromatin was cross-linked with 1% formaldehyde in PBS for 10 minutes at room temperature. After cross-linked cells were lysed and fragmented by sonication, chromatin fragments were immunoprecipitated with the antibodies to H3K4me3 (Millipore Sigma, Burlington, MA; 07-473), bound to Protein A/G PLUS-Agarose (Santa Cruz Biotechnology, SC2003). After extensive washing, the immunoprecipitated chromatin was eluted with 50 mM NaHCO_3_ 1% SDS, heated to 67 °C to reverse the cross-links, and the DNA purified by ethanol precipitation. Libraries were prepared using the DNA SMART ChIP-Seq Kit (Clonetech, Mountain View, CA). 10 ng of ChIP DNA was used as input for each sample. Libraries were sequenced on the Illumina NovaSeq6000 2×150bp. Reads were trimmed using Trim Galore (v0.6.7) before mapping to mm9 genome build using Bowtie (v2.4.1). Duplicates were removed and alignments cleaned for proper alignments pairs using Picard (v2.25.7) and Samtools (v1.9-4). Peaks were called using the broad option in MACS2 (v2.1.1), merged using BEDTools (v2.27) merge function, and overlapping reads within each library were quantified using BEDTools coverage function. Enrichment was compared between genotypes using EdgeR (v3.30.3) and plotting performed in R using psych (v2.1.6) and ComplexHeatmap (v2.4.3) packages. Data from Aldiri et al (GSE87064) was processed in same manner as above.

## DATA AVAILABILITY

ChIP-Seq data can be retrieved from the Sequence Read Archive (SRA) PRJNA778180.

## AUTHOR CONTRIBUTIONS

SC conceived of the study, CS and SC designed the experiments, CS and XZ performed experiments, CS analyzed data, PR performed bioinformatics analyses of ChIP-Seq results, CS, PR and SC wrote the manuscript. All authors read and approved the final manuscript.

## ACKNOWLEDGMENTS

Authors are grateful to Mingyan Yang, Guangyi Ling and Belinda Dana for technical assistance; Dr. Patricia Ernst and Dr. Francis Stewart for providing *Mll1 flox* and *Mll2 flox* mice, respectively. Authors thank the funding of NIH grants EY012543 (to SC) and EY002687 (to WU-DOVS), and unrestricted funds from Research to Prevent Blindness (to WU-DOVS)

## REFERENCES

Aldiri, I., Xu, B., Wang, L., Chen, X., Hiler, D., Griffiths, L., Valentine, M., Shirinifard, A., Thiagarajan, S., Sablauer, A., Barabas, M.-E., Zhang, J., Johnson, D., Frase, S., Zhou, X., Easton, J., Zhang, J., Mardis, E. R., Wilson, R. K., Downing, J. R., & Dyer, M. A. (2017). The Dynamic Epigenetic Landscape of the Retina During Development, Reprogramming, and Tumorigenesis. Neuron, 94(3), 550–568.e510.

Allis, C. D., Berger, S. L., Cote, J., Dent, S., Jenuwien, T., Kouzarides, T., Pillus, L., Reinberg, D., Shi, Y., Shiekhattar, R., Shilatifard, A., Workman, J., & Zhang, Y. (2007). New Nomenclature for Chromatin-Modifying Enzymes. Cell, 131(4), 633–636.

Applebury, M. L., Antoch, M. P., Baxter, L. C., Chun, L. L. Y., Falk, J. D., Farhangfar, F., Kage, K., Krzystolik, M. G., Lyass, L. A., & Robbins, J. T. (2000). The Murine Cone Photoreceptor: A Single Cone Type Expresses Both S and M Opsins with Retinal Spatial Patterning. Neuron, 27(3), 513–523.

Austin, C. P., & Cepko, C. L. (1995). Specification of Cell Fate in the Vertebrate Retina. In B. H. J. Juurlink, P. H. Krone, W. M. Kulyk, V. M. K. Verge, & J. R. Doucette (Eds.), Neural Cell Specification: Molecular Mechanisms and Neurotherapeutic Implications (pp. 139–143). Springer US.

Bassett, E. A., & Wallace, V. A. (2012). Cell fate determination in the vertebrate retina. Trends in Neurosciences, 35(9), 565–573.

Blanks, J. C., Adinolfi, A. M., & Lolley, R. N. (1974). Synaptogenesis in the photoreceptor terminal of the mouse retina. Journal of Comparative Neurology, 156(1), 81–93.

Brightman, D. S., Grant, R. L., Ruzycki, P. A., Suzuki, R., Hennig, A. K., & Chen, S. (2018). MLL1 is essential for retinal neurogenesis and horizontal inner neuron integrity. Scientific Reports, 8(1), 11902.

Brightman, D. S., Razafsky, D., Potter, C., Hodzic, D., & Chen, S. (2016). Nrl-Cre transgenic mouse mediates loxP recombination in developing rod photoreceptors. Genesis (New York, N.Y. : 2000), 54(3), 129–135.

Burmeister, M., Novak, J., Liang, M.-Y., Basu, S., Ploder, L., Hawes, N. L., Vidgen, D., Hoover, F., Goldman, D., Kalnins, V. I., Roderick, T. H., Taylor, B. A., Hankin, M. H., & McLnnes, R. R. (1996). Ocular retardation mouse caused by Chx10 homeobox null allele: impaired retinal progenitor proliferation and bipolar cell differentiation. Nature Genetics, 12(4), 376–384.

Cenik, B. K., & Shilatifard, A. (2021). COMPASS and SWI/SNF complexes in development and disease. Nature Reviews Genetics, 22(1), 38–58.

Cepko, C. (2014). Intrinsically different retinal progenitor cells produce specific types of progeny. Nature Reviews Neuroscience, 15(9), 615–627.

Cepko, C. L., Austin, C. P., Yang, X., Alexiades, M., & Ezzeddine, D. (1996). Cell fate determination in the vertebrate retina. Proceedings of the National Academy of Sciences of the United States of America, 93(2), 589–595.

Chen, S., Wang, Q. L., Nie, Z., Sun, H., Lennon, G., Copeland, N. G., Gilbert, D. J., Jenkins, N. A., & Zack, D. J. (1997). Crx, a novel Otx-like paired-homeodomain protein, binds to and transactivates photoreceptor cell-specific genes. Neuron, 19(5), 1017–1030.

Chen, Y., Anastassiadis, K., Kranz, A., Stewart, A. F., Arndt, K., Waskow, C., Yokoyama, A., Jones, K., Neff, T., Lee, Y., & Ernst, P. (2017). MLL2, Not MLL1, Plays a Major Role in Sustaining MLL-Rearranged Acute Myeloid Leukemia. Cancer Cell, 31(6), 755–770.e756.

Crump, N. T., & Milne, T. A. (2019). Why are so many MLL lysine methyltransferases required for normal mammalian development? Cellular and Molecular Life Sciences, 76(15), 2885–2898.

Denissov, S., Hofemeister, H., Marks, H., Kranz, A., Ciotta, G., Singh, S., Anastassiadis, K., Stunnenberg, H. G., & Stewart, A. F. (2014). Mll2 is required for H3K4 trimethylation on bivalent promoters in embryonic stem cells, whereas Mll1 is redundant. Development, 141(3), 526–537.

Dou, Y., Milne, T. A., Ruthenburg, A. J., Lee, S., Lee, J. W., Verdine, G. L., Allis, C. D., & Roeder, R. G. (2006). Regulation of MLL1 H3K4 methyltransferase activity by its core components. Nature Structural & Molecular Biology, 13(8), 713–719.

Dyer, M. A., Livesey, F. J., Cepko, C. L., & Oliver, G. (2003). Prox1 function controls progenitor cell proliferation and horizontal cell genesis in the mammalian retina. Nature Genetics, 34(1), 53–58.

Eissenberg, J. C., & Shilatifard, A. (2010). Histone H3 lysine 4 (H3K4) methylation in development and differentiation. Developmental Biology, 339(2), 240–249.

Emerson, Mark M., Surzenko, N., Goetz, Jillian J., Trimarchi, J., & Cepko, Constance L. (2013). Otx2 and Onecut1 Promote the Fates of Cone Photoreceptors and Horizontal Cells and Repress Rod Photoreceptors. Developmental Cell, 26(1), 59–72.

Furukawa, T., Morrow, E. M., & Cepko, C. L. (1997). Crx, a Novel otx-like Homeobox Gene, Shows Photoreceptor-Specific Expression and Regulates Photoreceptor Differentiation. Cell, 91(4), 531–541.

Furukawa, T., Ueno, A., & Omori, Y. (2020). Molecular mechanisms underlying selective synapse formation of vertebrate retinal photoreceptor cells. Cellular and Molecular Life Sciences, 77(7), 1251–1266.

Gan, T., Jude, C. D., Zaffuto, K., & Ernst, P. (2010). Developmentally induced Mll1 loss reveals defects in postnatal haematopoiesis. Leukemia, 24(10), 1732–1741.

Glaser, S., Schaft, J., Lubitz, S., Vintersten, K., van der Hoeven, F., Tufteland, K. R., Aasland, R., Anastassiadis, K., Ang, S.-L., & Stewart, A. F. (2006). Multiple epigenetic maintenance factors implicated by the loss of Mll2 in mouse development. Development, 133(8), 1423–1432.

Green, E. S., Stubbs, J. L., & Levine, E. M. (2003). Genetic rescue of cell number in a mouse model of microphthalmia:interactions between Chx10 and G1-phase cell cycle regulators. Development, 130(3), 539–552.

Gu, B., & Lee, M. G. (2013). Histone H3 lysine 4 methyltransferases and demethylases in self-renewal anddifferentiation of stem cells. Cell & Bioscience, 3(1), 39.

Guido, W. (2018). Development, form, and function of the mouse visual thalamus. Journal of Neurophysiology, 120(1), 211–225.

Hafler, B. P., Surzenko, N., Beier, K. T., Punzo, C., Trimarchi, J. M., Kong, J. H., & Cepko, C. L. (2012). Transcription factor Olig2 defines subpopulations of retinal progenitor cells biased toward specific cell fates. Proc Natl Acad Sci U S A, 109(20), 7882–7887.

Hennig, A. K., Peng, G. H., & Chen, S. (2008). Regulation of photoreceptor gene expression by Crx-associated transcription factor network. Brain Res, 1192, 114–133.

Jakovcevski, M., Ruan, H., Shen, E. Y., Dincer, A., Javidfar, B., Ma, Q., Peter, C. J., Cheung, I., Mitchell, A. C., Jiang, Y., Lin, C. L., Pothula, V., Stewart, A. F., Ernst, P., Yao, W.-D., & Akbarian, S. (2015). Neuronal Kmt2a/Mll1 Histone Methyltransferase Is Essential for Prefrontal Synaptic Plasticity and Working Memory. The Journal of Neuroscience, 35(13), 5097.

Jones, W. D., Dafou, D., McEntagart, M., Woollard, W. J., Elmslie, F. V., Holder-Espinasse, M., Irving, M., Saggar, A. K., Smithson, S., Trembath, R. C., Deshpande, C., & Simpson, M. A. (2012). De novo mutations in MLL cause Wiedemann-Steiner syndrome. American journal of human genetics, 91(2), 358–364.

Kerimoglu, C., Sakib, M. S., Jain, G., Benito, E., Burkhardt, S., Capece, V., Kaurani, L., Halder, R., Agís-Balboa, R. C., Stilling, R., Urbanke, H., Kranz, A., Stewart, A. F., & Fischer, A. (2017). KMT2A and KMT2B Mediate Memory Function by Affecting Distinct Genomic Regions. Cell Reports, 20(3), 538–548.

Kim, D. S., Ross, S. E., Trimarchi, J. M., Aach, J., Greenberg, M. E., & Cepko, C. L. (2008). Identification of molecular markers of bipolar cells in the murine retina. The Journal of comparative neurology, 507(5), 1795–1810.

Koike, C., Nishida, A., Ueno, S., Saito, H., Sanuki, R., Sato, S., Furukawa, A., Aizawa, S., Matsuo, I., Suzuki, N., Kondo, M., & Furukawa, T. (2007). Functional Roles of Otx2 Transcription Factor in Postnatal Mouse Retinal Development. Molecular and Cellular Biology, 27(23), 8318–8329.

Krivtsov, A. V., & Armstrong, S. A. (2007). MLL translocations, histone modifications and leukaemia stem-cell development. Nature Reviews Cancer, 7(11), 823–833.

Le, Y. Z., Ash, J. D., Al-Ubaidi, M. R., Chen, Y., Ma, J. X., & Anderson, R. E. (2004). Targeted expression of Cre recombinase to cone photoreceptors in transgenic mice. Mol Vis, 10, 1011–1018.

Li, B., Carey, M., & Workman, J. L. (2007). The Role of Chromatin during Transcription. Cell, 128(4), 707–719.

Liu, I. S., Chen, J. D., Ploder, L., Vidgen, D., van der Kooy, D., Kalnins, V. I., & McInnes, R. R. (1994). Developmental expression of a novel murine homeobox gene (Chx10): evidence for roles in determination of the neuroretina and inner nuclear layer. Neuron, 13(2), 377–393.

Livesey, F. J., & Cepko, C. L. (2001). Vertebrate neural cell-fate determination: Lessons from the retina. Nature Reviews Neuroscience, 2(2), 109–118.

Livne-bar, I., Pacal, M., Cheung, M. C., Hankin, M., Trogadis, J., Chen, D., Dorval, K. M., & Bremner, R. (2006). Chx10 is required to block photoreceptor differentiation but is dispensable for progenitor proliferation in the postnatal retina. Proceedings of the National Academy of Sciences of the United States of America, 103(13), 4988.

Lu, Y., Shiau, F., Yi, W., Lu, S., Wu, Q., Pearson, J. D., Kallman, A., Zhong, S., Hoang, T., Zuo, Z., Zhao, F., Zhang, M., Tsai, N., Zhuo, Y., He, S., Zhang, J., Stein-O’Brien, G. L., Sherman, T. D., Duan, X., Fertig, E. J., Goff, L. A., Zack, D. J., Handa, J. T., Xue, T., Bremner, R., Blackshaw, S., Wang, X., & Clark, B. S. (2020). Single-Cell Analysis of Human Retina Identifies Evolutionarily Conserved and Species-Specific Mechanisms Controlling Development. Developmental Cell, 53(4), 473–491.e479.

Mears, A. J., Kondo, M., Swain, P. K., Takada, Y., Bush, R. A., Saunders, T. L., Sieving, P. A., & Swaroop, A. (2001). Nrl is required for rod photoreceptor development. Nat Genet, 29.

Mellough, C. B., Bauer, R., Collin, J., Dorgau, B., Zerti, D., Dolan, D. W. P., Jones, C. M., Izuogu, O. G., Yu, M., Hallam, D., Steyn, J. S., White, K., Steel, D. H., Santibanez-Koref, M., Elliott, D. J., Jackson, M. S., Lindsay, S., Grellscheid, S., & Lako, M. (2019). An integrated transcriptional analysis of the developing human retina. Development, 146(2).

Meyers, R. M., Bryan, J. G., McFarland, J. M., Weir, B. A., Sizemore, A. E., Xu, H., Dharia, N. V., Montgomery, P. G., Cowley, G. S., Pantel, S., Goodale, A., Lee, Y., Ali, L. D., Jiang, G., Lubonja, R., Harrington, W. F., Strickland, M., Wu, T., Hawes, D. C., Zhivich, V. A., Wyatt, M. R., Kalani, Z., Chang, J. J., Okamoto, M., Stegmaier, K., Golub, T. R., Boehm, J. S., Vazquez, F., Root, D. E., Hahn, W. C., & Tsherniak, A. (2017). Computational correction of copy number effect improves specificity of CRISPR–Cas9 essentiality screens in cancer cells. Nature Genetics, 49(12), 1779–1784.

Milne, T. A., Briggs, S. D., Brock, H. W., Martin, M. E., Gibbs, D., Allis, C. D., & Hess, J. L. (2002). MLL Targets SET Domain Methyltransferase Activity to Hox Gene Promoters. Molecular Cell, 10(5), 1107–1117.

Morrow, E. M., Chen, C. M. A., & Cepko, C. L. (2008). Temporal order of bipolar cell genesis in the neural retina. Neural Development, 3(1), 2.

Nadal-Nicolás, F. M., Jiménez-López, M., Sobrado-Calvo, P., Nieto-López, L., Cánovas-Martínez, I., Salinas-Navarro, M., Vidal-Sanz, M., & Agudo, M. (2009). Brn3a as a Marker of Retinal Ganglion Cells: Qualitative and Quantitative Time Course Studies in Naïve and Optic Nerve– Injured Retinas. Investigative Ophthalmology & Visual Science, 50(8), 3860–3868.

Nishida, A., Furukawa, A., Koike, C., Tano, Y., Aizawa, S., Matsuo, I., & Furukawa, T. (2003). Otx2 homeobox gene controls retinal photoreceptor cell fate and pineal gland development. Nature Neuroscience, 6(12), 1255–1263.

Remez, Liv A., Onishi, A., Menuchin-Lasowski, Y., Biran, A., Blackshaw, S., Wahlin, K. J., Zack, D. J., & Ashery-Padan, R. (2017). Pax6 is essential for the generation of late-born retinal neurons and for inhibition of photoreceptor-fate during late stages of retinogenesis. Developmental Biology, 432(1), 140–150.

Roberts, M. R., Hendrickson, A., McGuire, C. R., & Reh, T. A. (2005). Retinoid X Receptor γ Is Necessary to Establish the S-opsin Gradient in Cone Photoreceptors of the Developing Mouse Retina. Investigative Ophthalmology & Visual Science, 46(8), 2897–2904.

Roberts, M. R., Srinivas, M., Forrest, D., Morreale de Escobar, G., & Reh, T. A. (2006). Making the gradient: Thyroid hormone regulates cone opsin expression in the developing mouse retina. Proceedings of the National Academy of Sciences, 103(16), 6218.

Rowan, S., & Cepko, C. L. (2004). Genetic analysis of the homeodomain transcription factor Chx10 in the retina using a novel multifunctional BAC transgenic mouse reporter. Developmental Biology, 271(2), 388–402.

Ruzycki, P. A., Zhang, X., & Chen, S. (2018). CRX directs photoreceptor differentiation by accelerating chromatin remodeling at specific target sites. Epigenetics & Chromatin, 11(1), 42.

Sapkota, D., Chintala, H., Wu, F., Fliesler, S. J., Hu, Z., & Mu, X. (2014). Onecut1 and Onecut2 redundantly regulate early retinal cell fates during development. Proceedings of the National Academy of Sciences, 111(39), E4086.

Shilatifard, A. (2008). Molecular implementation and physiological roles for histone H3 lysine 4 (H3K4) methylation. Current Opinion in Cell Biology, 20(3), 341–348.

Shinsky, S. A., Monteith, K. E., Viggiano, S., & Cosgrove, M. S. (2015). Biochemical Reconstitution and Phylogenetic Comparison of Human SET1 Family Core Complexes Involved in Histone Methylation*. Journal of Biological Chemistry, 290(10), 6361–6375.

Slany, R. K. (2016). The molecular mechanics of mixed lineage leukemia. Oncogene, 35(40), 5215–5223.

Soto, F., Ma, X., Cecil, J. L., Vo, B. Q., Culican, S. M., & Kerschensteiner, D. (2012). Spontaneous Activity Promotes Synapse Formation in a Cell-Type-Dependent Manner in the Developing Retina. The Journal of Neuroscience, 32(16), 5426.

Stenkamp, D. L. (2015). Development of the Vertebrate Eye and Retina. Progress in molecular biology and translational science, 134, 397–414.

Swaroop, A., Kim, D., & Forrest, D. (2010). Transcriptional regulation of photoreceptor development and homeostasis in the mammalian retina [Review Article]. Nature Reviews Neuroscience, 11, 563.

Szatko, K. P., Korympidou, M. M., Ran, Y., Berens, P., Dalkara, D., Schubert, T., Euler, T., & Franke, K. (2020). Neural circuits in the mouse retina support color vision in the upper visual field. Nature Communications, 11(1), 3481.

Sze, C. C., & Shilatifard, A. (2016). MLL3/MLL4/COMPASS Family on Epigenetic Regulation of Enhancer Function and Cancer. Cold Spring Harbor perspectives in medicine, 6(11), a026427.

Tian, N. (2004). Visual experience and maturation of retinal synaptic pathways. Vision Research, 44(28), 3307–3316.

Tran, N. M., Zhang, A., Zhang, X., Huecker, J. B., Hennig, A. K., & Chen, S. (2014). Mechanistically distinct mouse models for CRX-associated retinopathy. PLoS genetics, 10(2), e1004111–e1004111.

Turner, D. L., Snyder, E. Y., & Cepko, C. L. (1990). Lineage-independent determination of cell type in the embryonic mouse retina. Neuron, 4(6), 833–845.

Wu, F., Li, R., Umino, Y., Kaczynski, T. J., Sapkota, D., Li, S., Xiang, M., Fliesler, S. J., Sherry, D. M., Gannon, M., Solessio, E., & Mu, X. (2013). Onecut1 Is Essential for Horizontal Cell Genesis and Retinal Integrity. The Journal of Neuroscience, 33(32), 13053.

Wu, S., Chang, K.-C., & Goldberg, J. L. (2018). Retinal Cell Fate Specification. Trends in Neurosciences, 41(4), 165–167.

Yokoyama, A., Wang, Z., Wysocka, J., Sanyal, M., Aufiero Deborah, J., Kitabayashi, I., Herr, W., & Cleary Michael, L. (2004). Leukemia Proto-Oncoprotein MLL Forms a SET1-Like Histone Methyltransferase Complex with Menin To Regulate Hox Gene Expression. Molecular and Cellular Biology, 24(13), 5639–5649.

Young, R. W. (1985). Cell differentiation in the retina of the mouse. The Anatomical Record, 212(2), 199–205.

Yu, B. D., Hess, J. L., Horning, S. E., Brown, G. A. J., & Korsmeyer, S. J. (1995). Altered Hox expression and segmental identity in Mll-mutant mice. Nature, 378(6556), 505–508.

Zhang, T., Cooper, S., & Brockdorff, N. (2015). The interplay of histone modifications – writers that read. EMBO reports, 16(11), 1467–1481.

