## Supplementary Materials for "Essential functions of MLL1 and MLL2 in retinal development and cone cell maintenance"

Chi Sun[1], Xiaodong Zhang[1], Philip A. Ruzycki[1] and Shiming Chen[1,2]*
[1]Department of Ophthalmology and Visual Sciences, [2]Department of Developmental Biology, Washington University, Saint Louis, Missouri, USA

*To whom the correspondence should be addressed.
Mailing address:
660 South Euclid Avenue, MSC-8096-06-06,
St. Louis, MO 63110, USA


**Supplemental Materials**

### Supplemental Figures

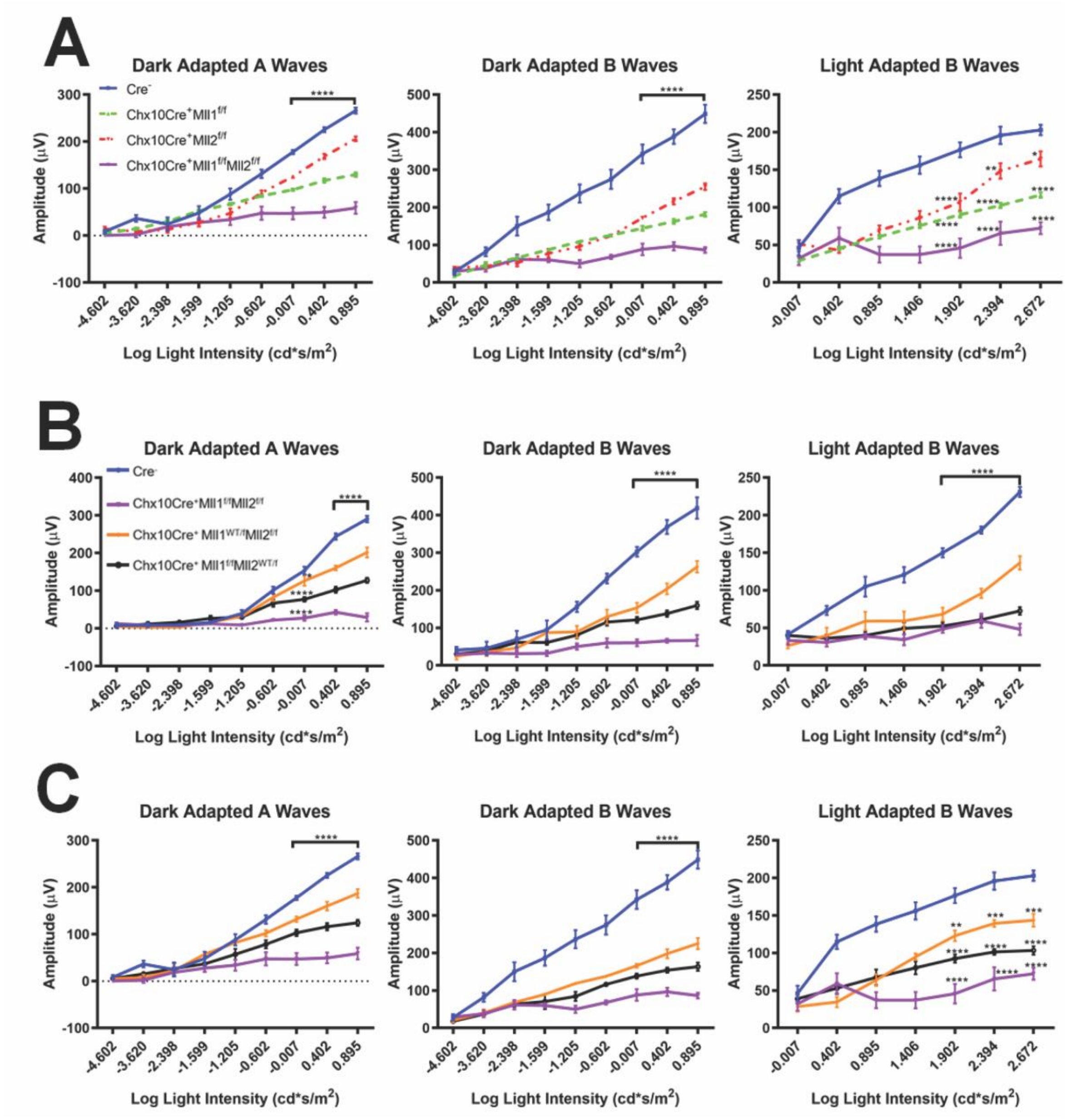


**Supplemental Figure 1**. (**A**) Dark adapted A-wave (left), dark adapted B-wave (center), light adapted B-wave (right) ERG of 2MO $Chx10Cre^{+}Mll1^{f/f}$, $Chx10Cre^{+}Mll2^{f/f}$, $Chx10Cre^{+}Mll1^{f/f}Mll2^{f/f}$, and $Chx10Cre^{-}$ mice. (**B**) Dark adapted A-wave (left), dark adapted B-wave (center), light adapted B-wave (right) ERG of 1MO $Chx10Cre^{+}Mll1^{WT/f}Mll2^{f/f}$, $Chx10Cre^{+}Mll1^{f/f}Mll2^{WT/f}$, $Chx10Cre^{+}Mll1^{f/f}Mll2^{f/f}$, and $Chx10Cre^{-}$ mice. (**C**) Dark adapted A-wave (left), dark adapted B-wave (center), light adapted B-wave (right) ERG analysis of 2MO $Chx10Cre^{+}Mll1^{WT/f}Mll2^{f/f}$, $Chx10Cre^{+}Mll1^{f/f}Mll2^{WT/f}$, $Chx10Cre^{+}Mll1^{f/f}Mll2^{f/f}$, and $Chx10Cre^{-}$ mice. Mean amplitudes (μV) are plotted against stimulus light intensity. Error bars represent SEM ($n \geq 4$). All statistics is done by two-way ANOVA and Tukey's multiple comparisons. Asterisks (*, **, ***, ****) denote $p \leq 0.05$, $p \leq 0.01$, $p \leq 0.001$, and $p \leq 0.0001$, respectively.

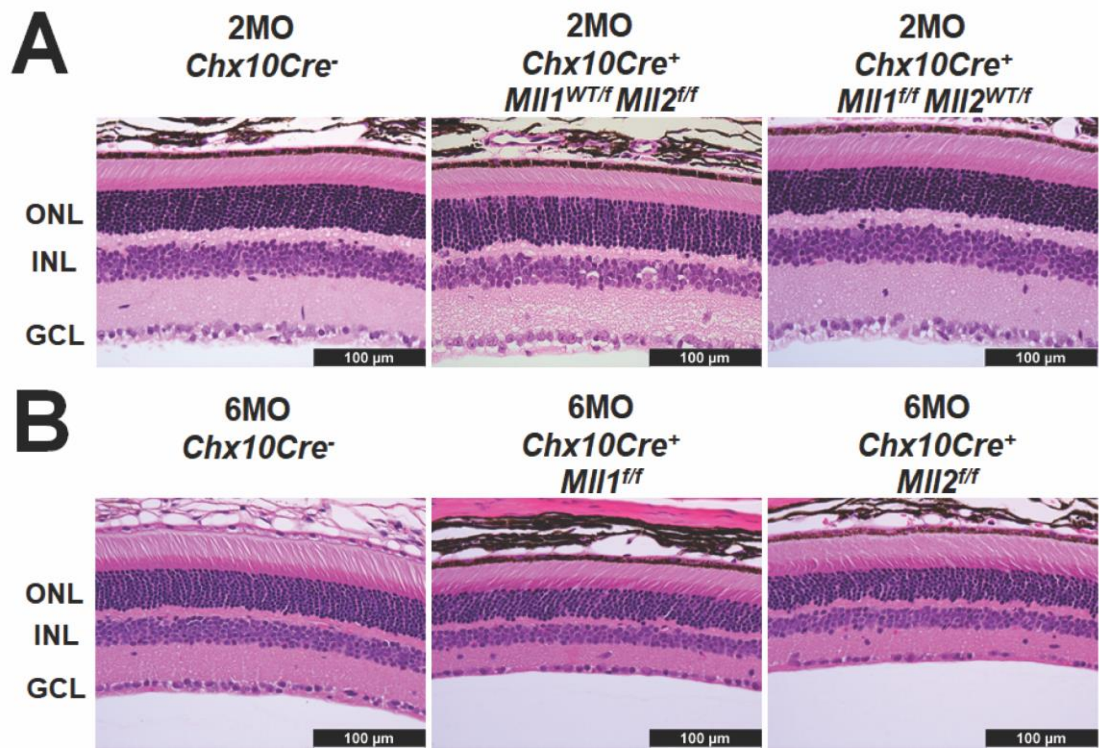


**Supplemental Figure 2.** Representative images of H & E-stained retinal cross-sections of the mice of indicated genotypes at 2MO (**A**) and 6MO (**B**). Scale bar = 100μm for all image panels.

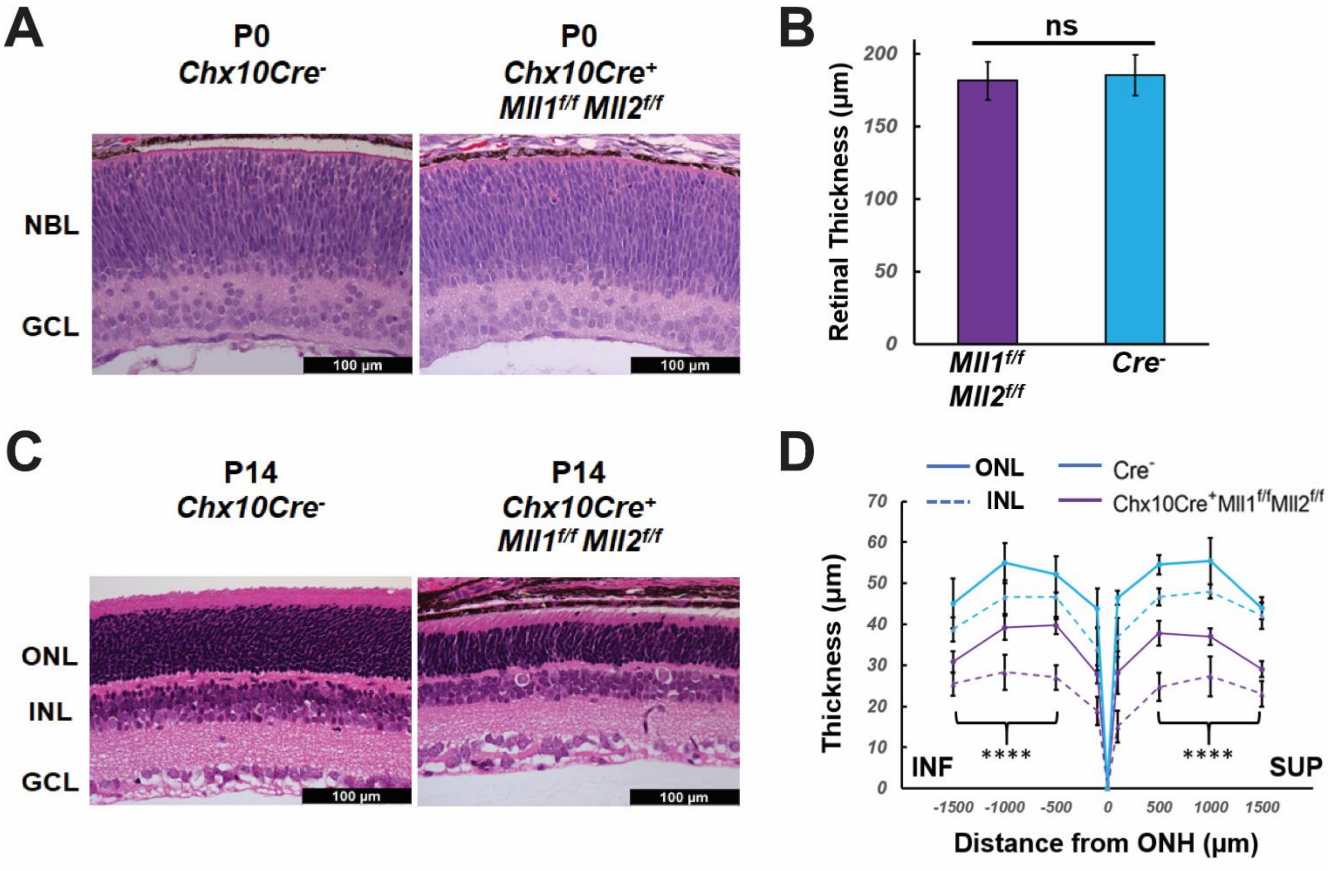


**Supplemental Figure 3.** (**A**) H & E -stained retinal cross-sections of P0 *Chx10Cre*$^{-}$ and *Chx10Cre*$^{+}$*Mll1*$^{f/f}$*Mll2*$^{f/f}$ retinae. (**B**) Retinal thickness (μm) in P0 retinae at 500μm from the ONH. All statistics is done by Tukey's multiple comparisons. Asterisks (****) denote $P \leq 0.0001$, ns means not significant. (**C**) H & E -stained retinal cross-sections of P14 *Chx10Cre*$^{-}$ and *Chx10Cre*$^{+}$*Mll1*$^{f/f}$*Mll2*$^{f/f}$ retinae. Scale bar = 100μm for all image panels. (**D**) Plots of ONL (solid line) and INL (dotted line) thickness (μm) for P14 retinae at the indicated positions from the ONH. SUP and INF indicate superior and inferior sides of the retina. Error bars represent SEM (n≥4). Results of statistical analysis for both ONL and INL thickness are shown.

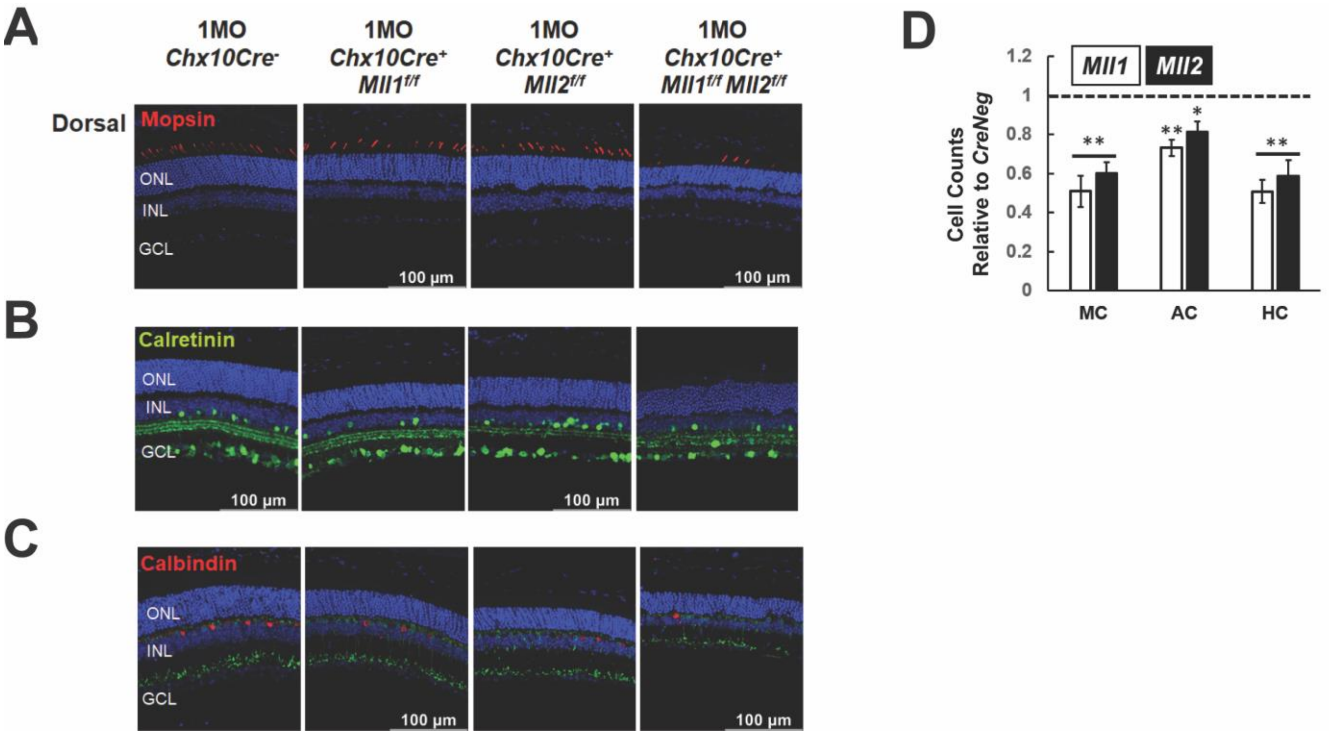


**Supplementary Figure 4**. Immunostaining of the following cellular markers in 1MO retinae of the indicated genotypes: Medium-wave-sensitive opsin 1 (Mopsin, red in **A**), Calretinin (green in **B**) for amacrine and ganglion cells, Calbindin D-28 (red in **C**) for horizontal cells. Nuclei are counterstained by DAPI (blue). Scale bar = 100μm for all image panels. (**D**) Cell counts for different retinal cell types in 1MO *Chx10Cre*$^{+}$*Mll1*$^{f/f}$ and *Chx10Cre*$^{+}$*Mll2*$^{f/f}$ retinae, normalized to counts from *Chx10Cre*$^{-}$ (*CreNeg*) littermates. Error bars represent SEM (n≥3). Asterisks (*, **) denote $p \leq 0.05$ and $p \leq 0.01$ respectively, ns means not significant by one-way ANOVA. MC=M cone photoreceptors, AC=amacrine cells, HC=horizontal cells.

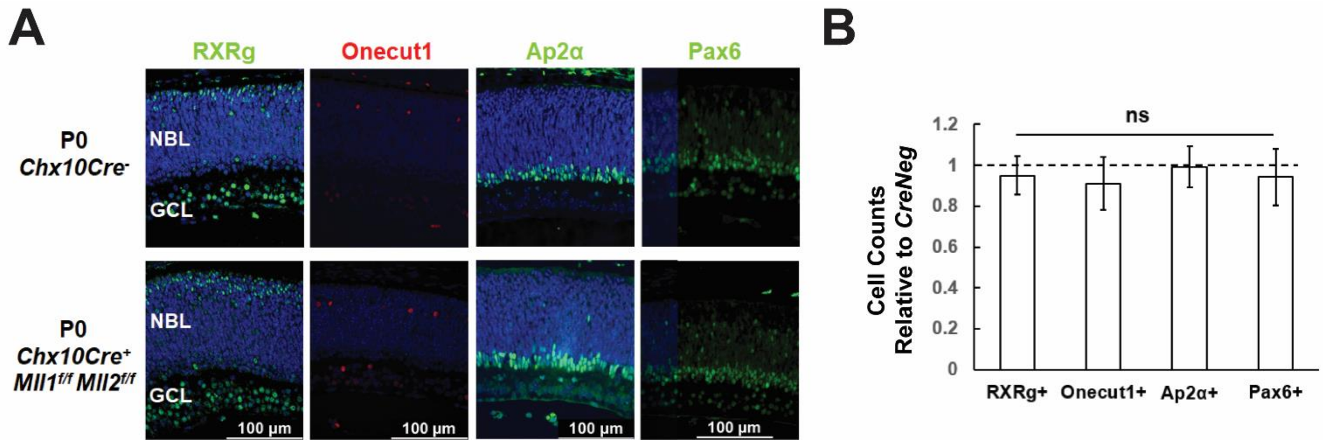


**Supplemental Figure 5. (A)** Immunostaining of the following cellular markers in P0 retinae of the indicated genotypes: (**A**) RXRg for fated cones (green in 1$^{st}$ panel from left), Onecut1 for fated horizontal cells (red in 2$^{nd}$ panel), Ap2α for early-born amacrine cells (green in 3$^{rd}$ panel), Pax6 for late-born progenitor cells (green in 4$^{th}$ panel). Nuclei are counterstained by DAPI (blue). Scale bar = 100µm for all image panels. (**B**) Cell counts for different cell types in P0 *Chx10Cre*$^{+}$*Mll1*$^{f/f}$*Mll2*$^{f/f}$ retinae, normalized to counts from *Chx10Cre*$^{-}$ (*CreNeg*) littermates. Error bars represent SEM (n≥3). ns means not significant by T-test.

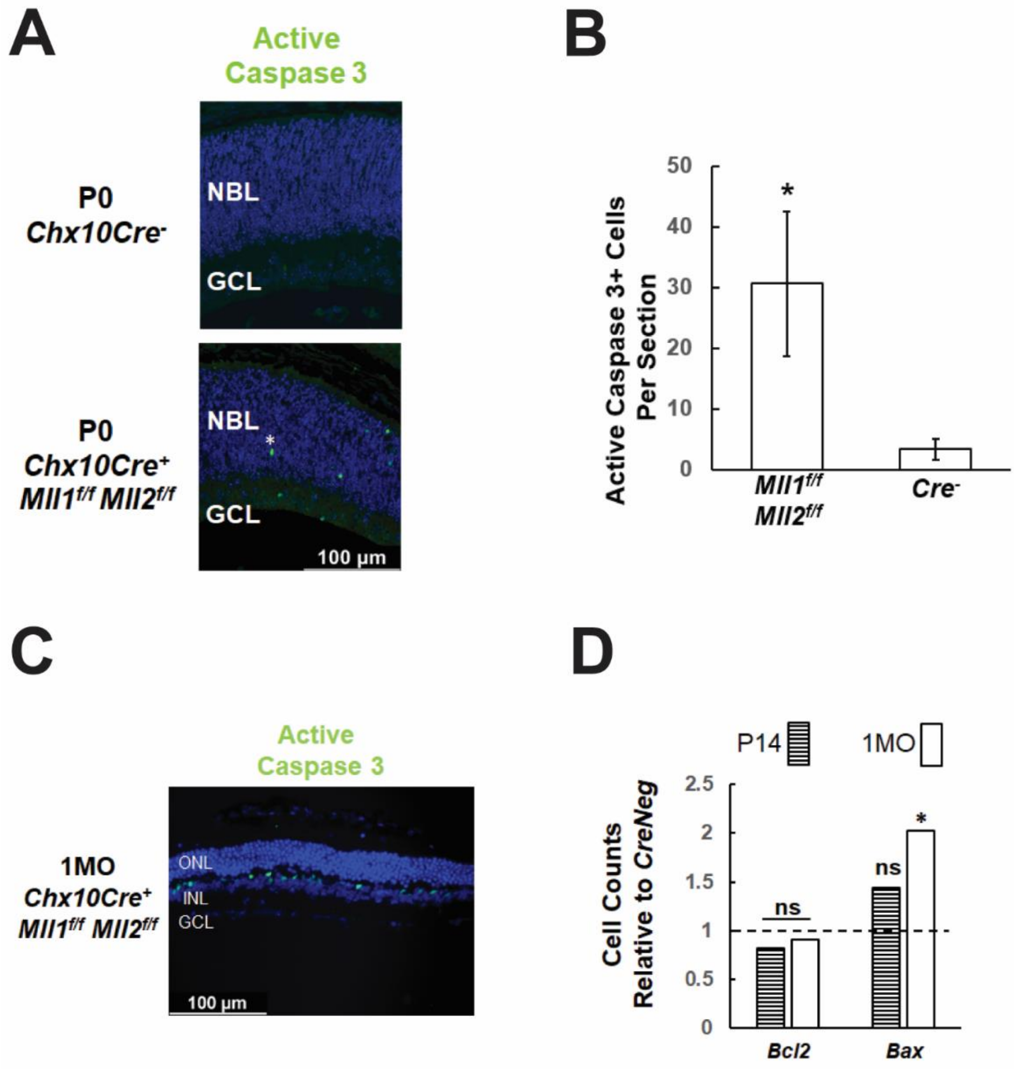


**Supplemental Figure 6.** (**A**) Active caspase 3 (green) immunostaining in P0 *Chx10Cre*$^-$ and *Chx10Cre*$^+$*Mll1*$^{f/f}$*Mll2*$^{f/f}$ retinae. Active caspase 3 immunoreactivity labels cells undergoing apoptosis. (**B**) Cell counts for Active caspase 3+ cells in P0 retinae. (**C**) Active caspase 3 (green) immunostaining in 1MO *Chx10Cre*$^+$*Mll1*$^{f/f}$*Mll2*$^{f/f}$ retinae. Nuclei are counterstained by DAPI (blue). Scale bar = 100µm for all image panels. (**D**) qRT-PCR analysis of *Bax* and *Bcl2* in P14 and 1MO *Chx10Cre*$^+$*Mll1*$^{f/f}$*Mll2*$^{f/f}$ retinae. Results are plotted as relative expression to *Chx10Cre*$^-$ (*CreNeg*) littermate controls (n≥4). Asterisks (*) denote $p \leq 0.05$ by one-way ANOVA, and ns means not significant.

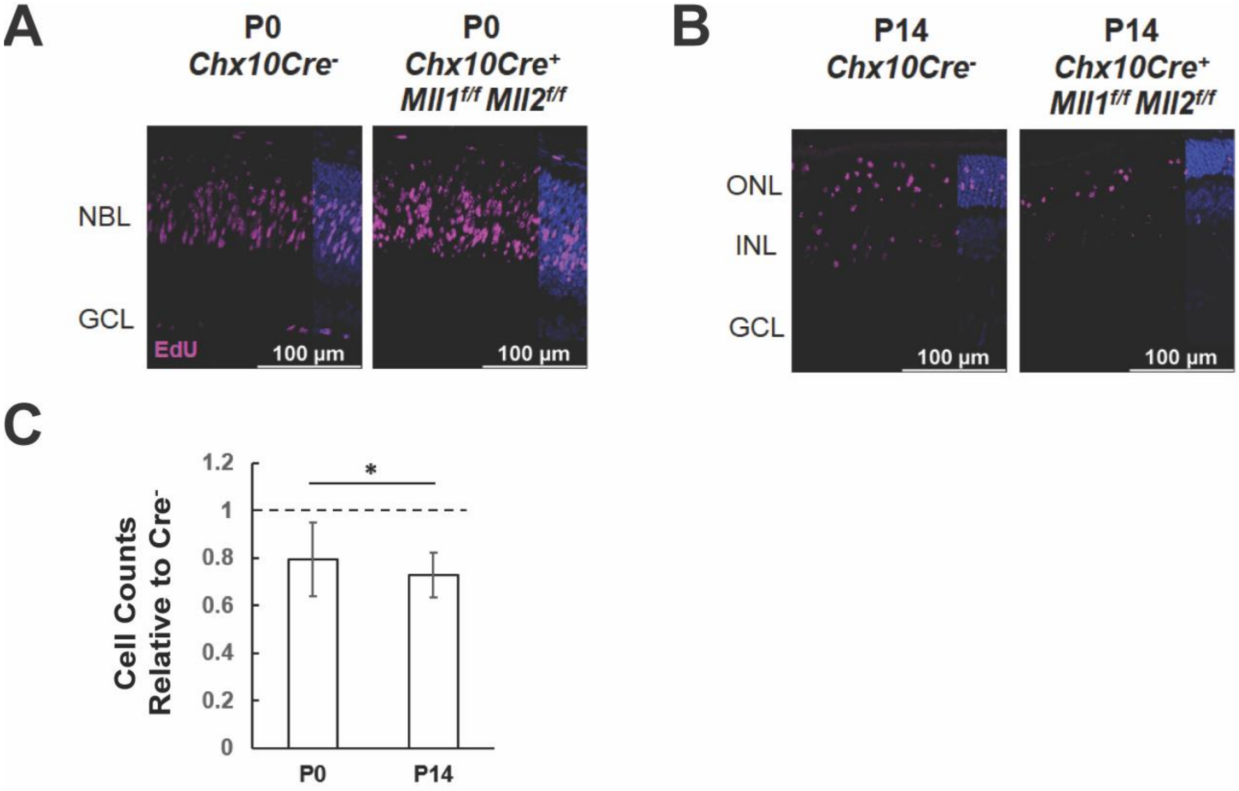


**Supplemental Figure 7.** EdU immunostaining (red) on P0 (**A**) and P14 (**B**) retinal cross-sections of *Chx10Cre*$^-$ and *Chx10Cre*$^+$*Mll1*$^{f/f}$*Mll2*$^{f/f}$ retinae. Nuclei are counterstained by DAPI (blue). Scale bar = 100µm for all image panels. (**C**) Cell counts for EdU+ cells in P0 and P14 *Chx10Cre*$^+$*Mll1*$^{f/f}$*Mll2*$^{f/f}$ retinae, normalized to counts from *Chx10Cre*$^-$ littermates. Error bars represent SEM (n≥3). Asterisks (*) denote $p \leq 0.05$ by one-way ANOVA.

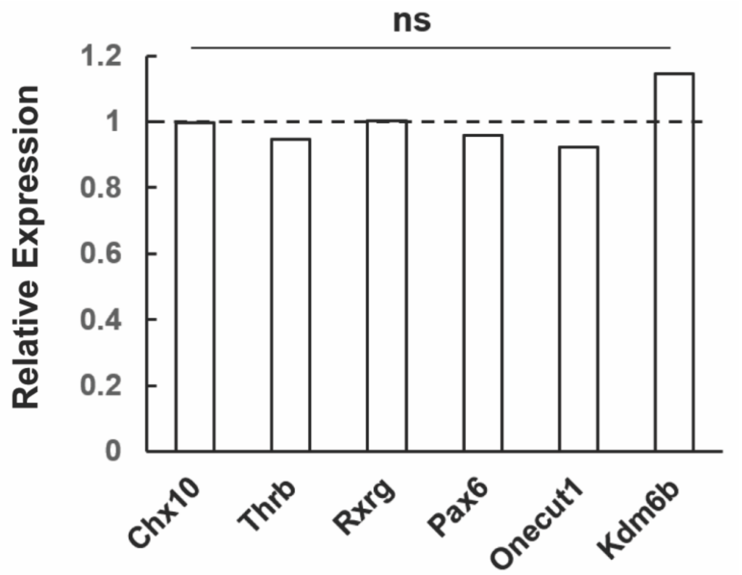


**Supplemental Figure 8.** qRT-PCR analysis of selected genes in P0 *Chx10Cre*$^{+}$*Mll1*$^{f/f}$*Mll2*$^{f/f}$ retinae. Results are plotted as relative expression to *Chx10Cre*$^{-}$ littermate controls (n≥4). ns means not significant by T-test.

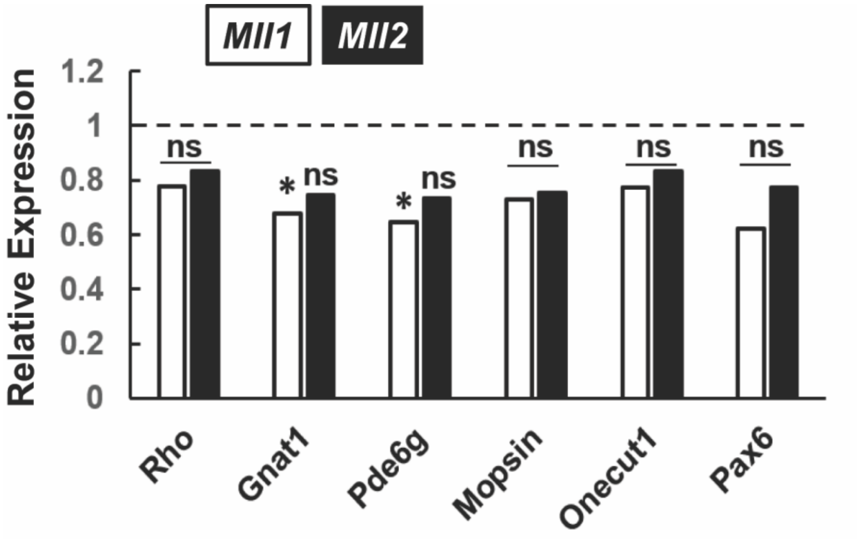


**Supplemental Figure 9.** qRT-PCR analysis of selected genes in 1MO *Chx10Cre*$^{+}$*Mll1*$^{f/f}$ and *Chx10Cre*$^{+}$*Mll2*$^{f/f}$ retinae. Results are plotted as relative expression to *Chx10Cre*$^{-}$ littermate controls (n≥4). Asterisks (*) denote P ≤ 0.05 by T-test, ns means not significant.

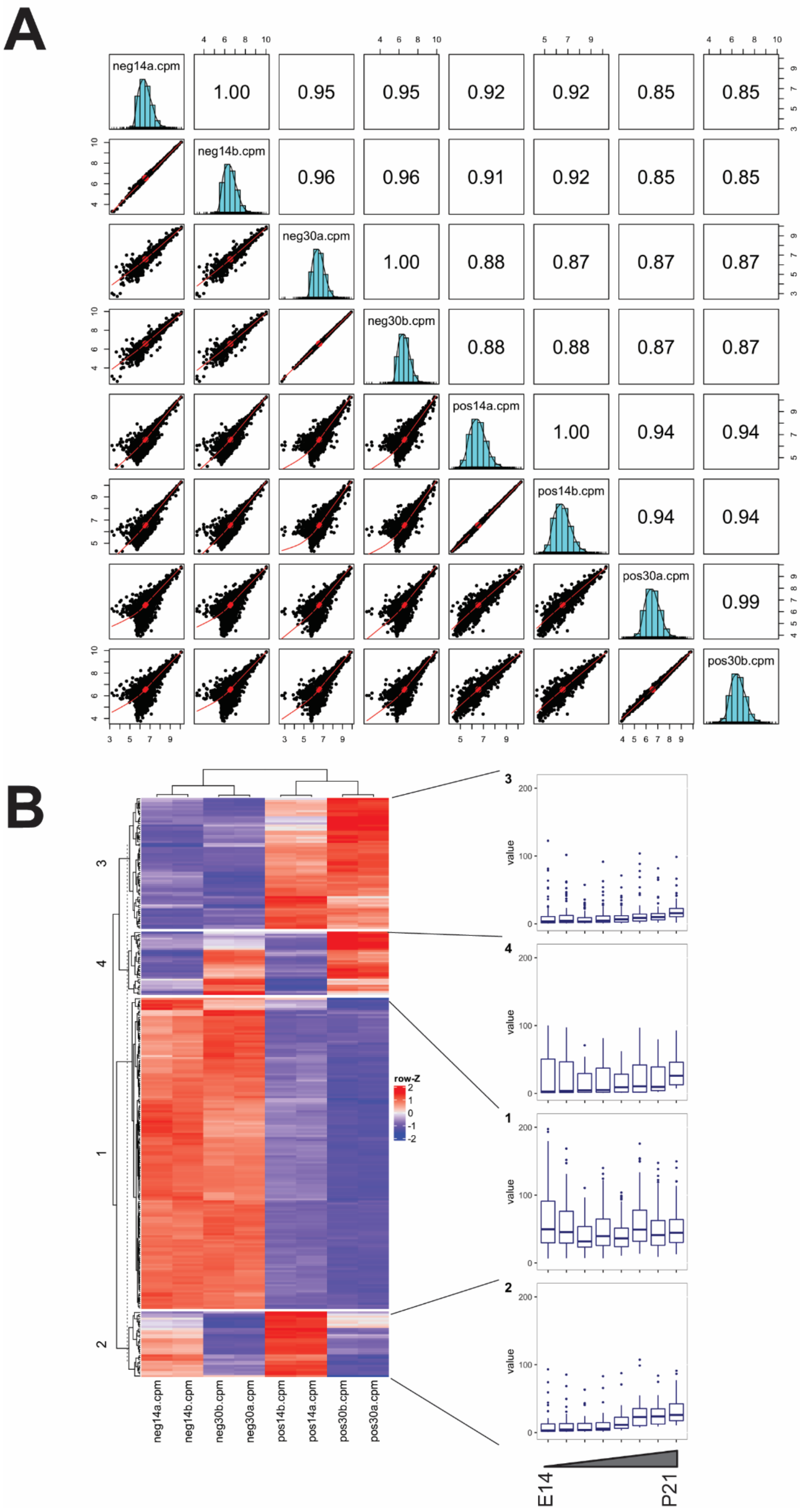
A
neg14a.cpm
neg14b.cpm
neg30a.cpm
neg30b.cpm
pos14a.cpm
pos14b.cpm
pos30a.cpm
pos30b.cpm
1.00
0.95
0.95
0.92
0.92
0.85
0.85
0.96
0.96
0.91
0.92
0.85
0.85
1.00
0.88
0.87
0.87
0.87
0.88
0.88
0.87
0.87
1.00
0.94
0.94
0.94
0.94
0.99
B
row-Z
3
4
1
2
neg14a.cpm
neg14b.cpm
neg30b.cpm
neg30a.cpm
pos14b.cpm
pos14a.cpm
pos30b.cpm
pos30a.cpm
value
E14
P21

**Supplemental Figure 10.** (**A**) Correlation plot comparing each library by scatterplot and Pearson correlation value. (**B**) Clustered heatmap of all 279 H3K4me3 peaks that showed differential deposition in pairwise comparisons between all samples (>2 fold-change, $P < 0.05$). Boxplots display enrichment of indicated histones profiled over retinal development (E14-P21) in Aldiri et al., 2017.

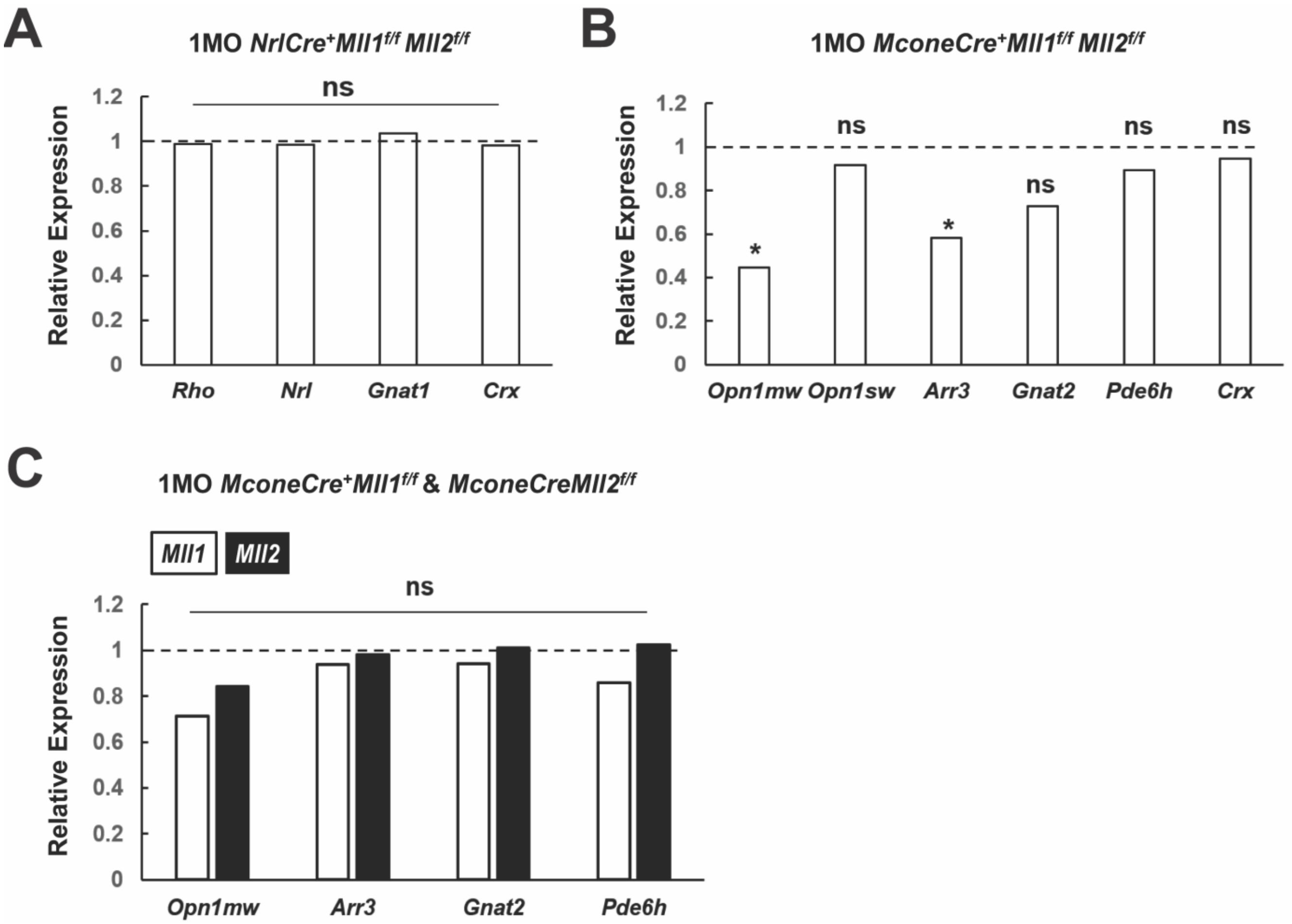


**Supplemental Figure 11.** (**A**) qRT-PCR analysis of selected genes in 1MO *NrlCre*$^{+}$*Mll1*$^{f/f}$*Mll2*$^{f/f}$ retinae. Results are plotted as relative expression to *NrlCre*$^{-}$ littermate controls (n≥4). (**B**) qRT-PCR analysis of selected genes in 1MO *MconeCre*$^{+}$*Mll1*$^{f/f}$*Mll2*$^{f/f}$ retinae. Results are plotted as relative expression to *MconeCre*$^{-}$ littermate controls (n≥4). (**C**) qRT-PCR analysis of selected genes in 1MO *MconeCre*$^{+}$*Mll1*$^{f/f}$ and *MconeCre*$^{+}$*Mll2*$^{f/f}$ retinae. Results are plotted as relative expression to *MconeCre*$^{-}$ littermate controls (n≥4). Asterisks (*) denote $p \leq 0.05$ by T-test, ns means not significant.

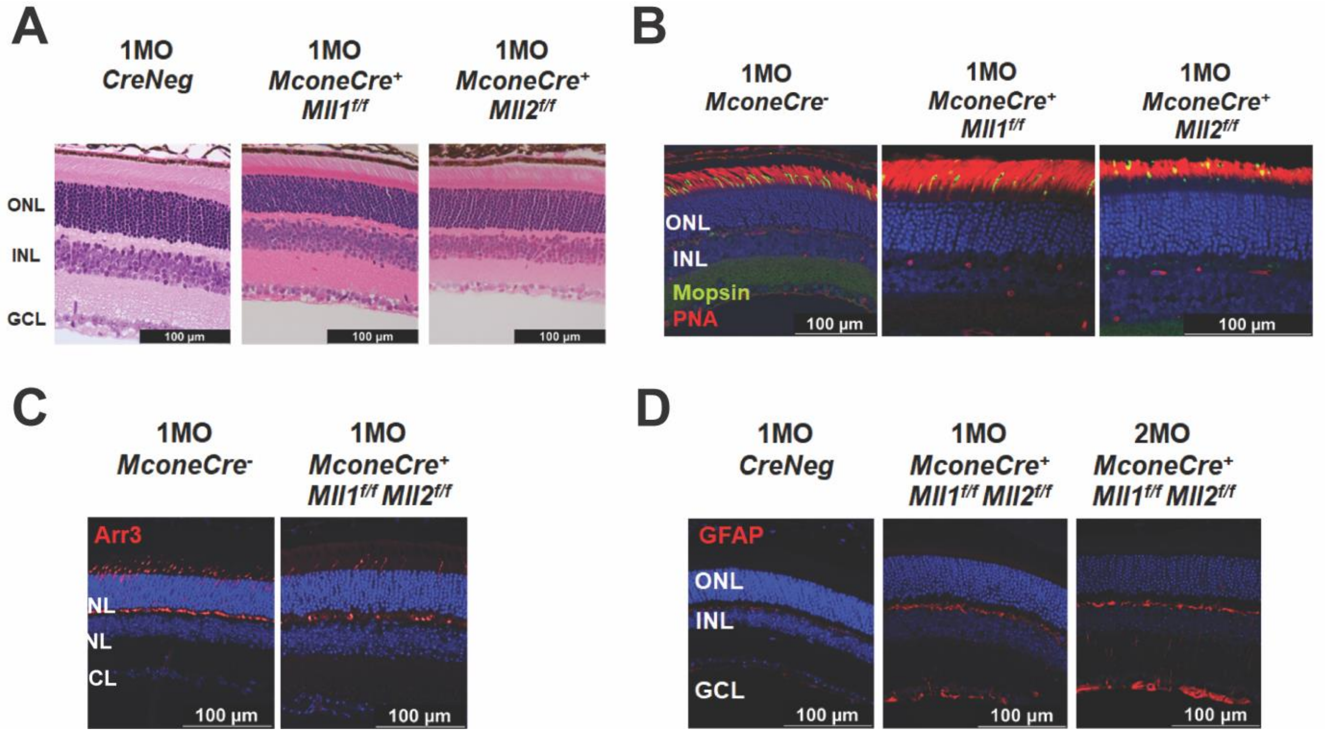


**Supplemental Figure 12.** (**A**) H & E-stained retinal cross-section of 1MO *Cre*$^{-}$ (*CreNeg*), *MconeCre*$^{+}$*Mll1*$^{f/f}$, *MconeCre*$^{+}$*Mll2*$^{f/f}$ retinae. (**B**) Mopsin and PNA immunostaining in 1MO *MconeCre*$^{-}$, *MconeCre*$^{+}$*Mll1*$^{f/f}$, and *MconeCre*$^{+}$*Mll2*$^{f/f}$ retinae. (**C**) Cone arrestin (Arr3) immunostaining in 1MO *MconeCre*$^{-}$ and *MconeCre*$^{+}$*Mll1*$^{f/f}$*Mll2*$^{f/f}$ retinae. (**D**) GFAP immunostaining in 1MO *MconeCre*$^{-}$ (*CreNeg*), 1MO and 2MO *MconeCre*$^{+}$*Mll1*$^{f/f}$*Mll2*$^{f/f}$ retinae. Nuclei are counterstained by DAPI (blue). Scale bar = 100µm for all image panels.

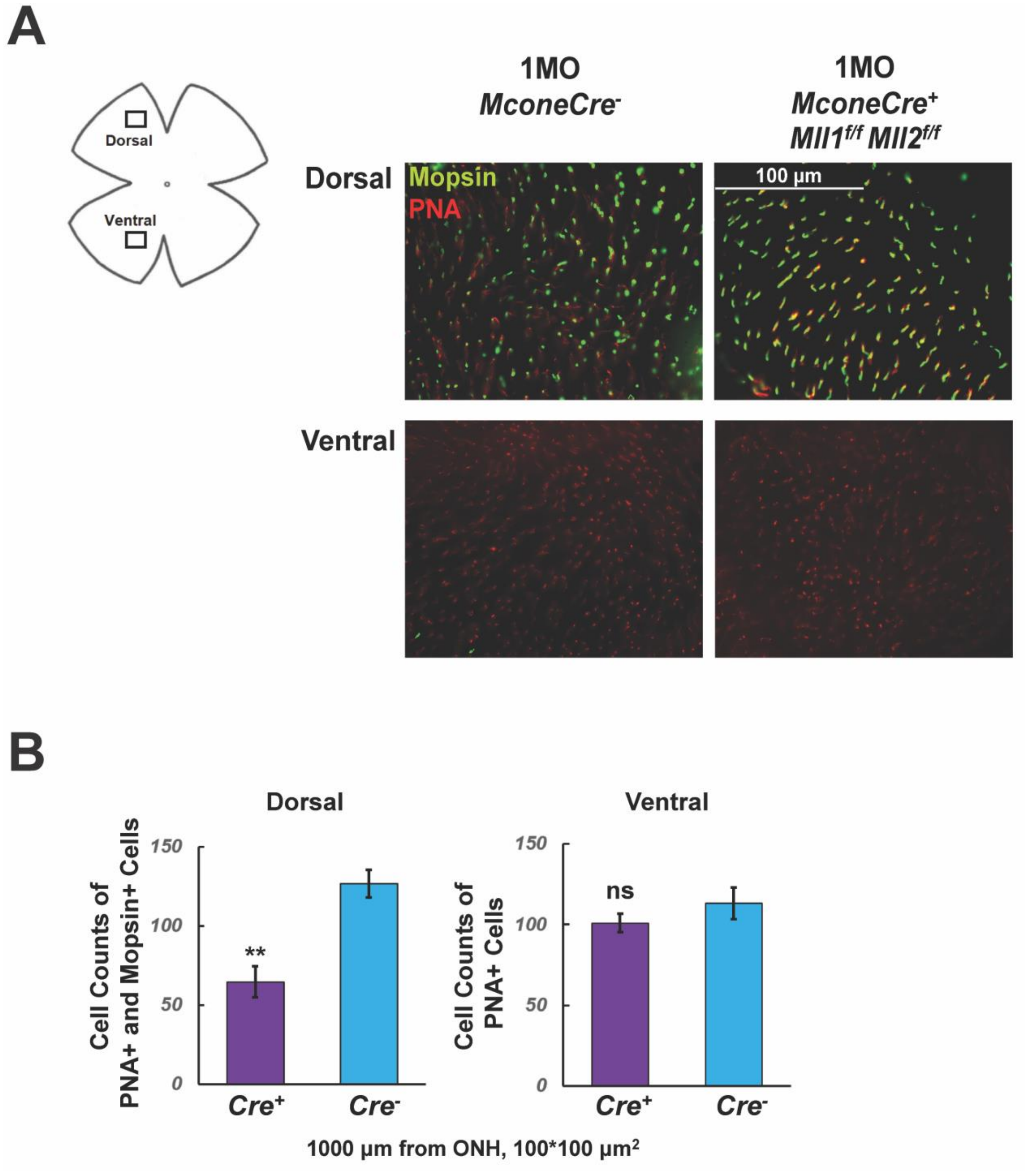


**Supplemental Figure 13.** (**A**) Whole-amount images of Mopsin (green) and PNA (red) immunostaining in 1MO *MconeCre*$^{-}$ and *MconeCre*$^{+}$*Mll1*$^{f/f}$*Mll2*$^{f/f}$ retinae. Top panel contains images taken at the dorsal retinae; bottom panel contains images taken at the ventral retinae. (**B**) Cell counts for Mopsin and PNA co-labelled cells at dorsal (left) and ventral (right) in 1MO *MconeCre*$^{-}$ and *MconeCre*$^{+}$*Mll1*$^{f/f}$*Mll2*$^{f/f}$ retinae. Error bars represent SEM (n=3). Cell counting is taken within an area of 100*100 µm$^2$, 1000 µm from ONH. Asterisks (**) denote $p \leq 0.01$ by T-test, ns means not significant.

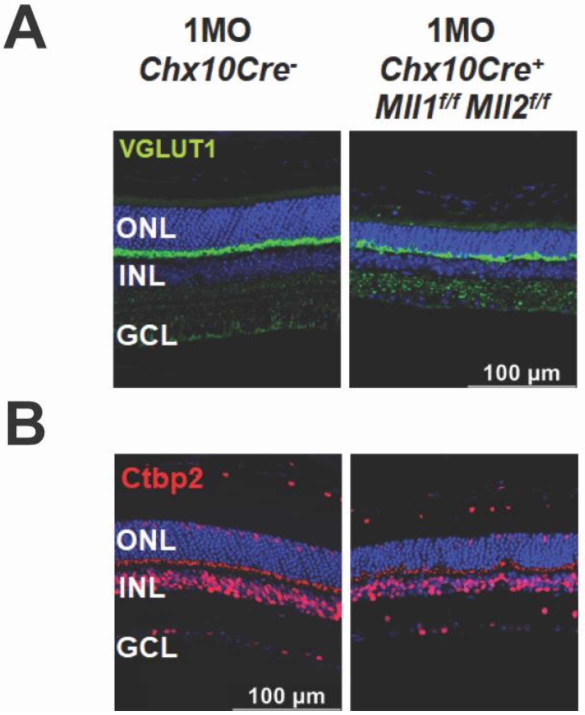


**Supplemental Figure 14.** Immunostaining of the following biomarkers in 1MO retinae of the indicated genotypes: (**A**) Vesicular glutamate transporter 1 (VGLUT1 in gren) and (**B**) C-terminal binding protein 2 (Ctbp2 in red) Nuclei are counterstained by DAPI (blue). Scale bar = 100µm for all image panels. VGLUT1 immunoreactivity labels the membrane of synaptic vesicles, Ctbp2/RIBEYE immunoreactivity labels ribbon synapses.

### Supplemental Tables

#### Supplemental Table 1: Antibodies

| Antibodies | Source | Species |
|---|---|---|
| Active Caspase-3 | R&D Systems AF835 | Rabbit |
| Ap2α | DSHB 3B5 | Mouse |
| Brn3a | MilliporeSigma MAB1585 | Mouse |
| Calbindin D28K | MilliporeSigma C9848 | Mouse |
| Calretinin | MilliporeSigma AB5054 | Rabbit |
| Cone Arrestin | MilliporeSigma AB15282 | Rabbit |
| Ctbp2 | BD Biosciences 612044 | Mouse |
| GFAP | Agilent z0334 | Rabbit |
| Glutamine Synthetase | BD Biosciences 610517 | Mouse |
| Iba1 | WakoFujifilm 019-19741 | Rabbit |
| Ki67 | BD Biosciences 550809 | Mouse |
| M-opsin (Opn1mw) | MilliporeSigma AB5405 | Rabbit |
| Onecut1 | Santa Cruz sc13050 | Rabbit |
| Pax6 | DSHB Pax6 | Mouse |
| Phospho-Histone H3 | MilliporeSigma 06-570 | Rabbit |
| PKCA | MilliporeSigma P5704 | Mouse |
| PNA-Rhodamine | Vector Labs RL-1072 | lectins |
| Rhodopsin (RetP1) | MilliporeSigma O4886 | Mouse |
| S-opsin (Opn1sw) | Santa Cruz sc-14363 | Goat |
| RXRg | Santa Cruz sc-555 | Rabbit |
| VGluT1 | MilliporeSigma AB5905 | Guinea pig |

**Supplemental Table 2: Primer Sets for qPCR**

| Gene | Forward Primer (5' – 3') | Reverse Primer (5' – 3') |
|---|---|---|
| *Actb* | CCAACTGGGACGACATGGAG | TGGTACGACCAGAGGCATACA |
| *Arr3* | AAGTTTTCCATCTACCTGGGG | TCACATCCAAGTCATCACGG |
| *Bax* | GCAGAGGATGATTGCTGACG | TTCCAGATGGTGAGCGAGG |
| *Bcl2* | AAAATACAGCATTGCGGAGG | TCCAGCATCCCACTCGTAGC |
| *Chx10* | CTCCCAGAAGACAGGATACAGGTG | TGCCTCCAGCGACTTTTTGTG |
| *Crx* | TGTCCCATACTCAAGTGCCC | TGCTGTTTCTGCTGCTGTCG |
| *Ctbp2* | TGGTGGATGAGAAAGCCTTAGC | ATGCGACCTGTGATTGCTCG |
| *Gfap* | GAAGAAAACCGCATCACCATTC | CCAGAAGGAAGGGAAGTGCTG |
| *Gnat1* | ACGATGGACCTAACACTTACGAGG | TGGAAAGGACGGTATTTGAGG |
| *Gnat2* | AGTCAAGACAACAGGCATCATCG | TCACTTCGTCATCCTCCACCAG |
| *Grm6* | CCGCATCTACCGCATTTTCG | TCCACCGTCCTCTGTTCCTCATA |
| *Kdm6b* | GTCTGATGCCAAGAGGTGGAAG | GCTGATGGTCTCCCAATAGTGC |
| *Mll1* | TGAGTACAACCCTAACGATGAGGAA | CGGAATCTCATGGGCATTG |
| *Mll2* | AAAGACATCCAAAGAGGCTGTGG | TGTAGCACCCAATACCCTTCCC |
| *Nr2e3* | AGTCCCAGGTGATGCTAAGC | TTCTAAGATGTGCTGCCCC |
| *Nrl* | TTCTGGTTCTGACAGTGACTACG | AAGGCTCCCGCTTTATTTC |
| *Onecut1* | GGAAAGAGCAAGAACACGG | GATGAGGACGATGAACTGC |
| *Onecut2* | GATGTCTCACCTCAATGGC | TGGTTCTTGCTCTTTGCG |
| *Opn1mw* | GGTGGTGATGGTCTTCGCATAC | TTGGAGGTGCTGGAAAGTTCAG |
| *Opn1sw* | GCTGGACTTACGGCTTGTCACC | TGTGGCGTTGTGTTTGCTGC |
| *Pax6* | CCAGTGTCTACCAGCCAATCCC | GGTGAAATGAGTCCTGTTGAAGTGG |
| *Pkca* | CATTGCCCCAGAGATAATCGC | GGTGTTTGGTCATAAGTCCTTTGC |
| *Pde6a* | CACTCCTGAGAGATGAGAGCC | CAGGGTTTGGTGATGGCTG |
| *Pde6b* | CAAGAAAGTGGGCACAGAAG | ATAGGCAGAGTCCGTATGC |
| *Pde6g* | GCAAGGGTTTGGGGATGACA | CGTGCAGCTCTAGGTGATTGAA |
| *Pde6h* | ACGGGAACACATTCGGCTC | CCAGATGGGTGAACGCTTC |
| *Prox1* | CAGAAGGACTCTCTTTGTCAC | GCTGAACCACTTGATGAGC |
| *Rho* | GCTTCCCTACGCCAGTGTG | CAGTGGATTCTTGCCGCAG |
| *RXRg* | CGTGGAGAACTCAACAAATGACCC | TGGATAAACCCTTGGCATCTGG |
| *Thrb* | CAACCTGGATGACACTGAAGTCG | TCTAAGAACAGAGGCGGGAAGAG |
| *Ubb* | CAACATCCAGAAAGAGTCAACC | ATGTTGTAATCAGAGAGGGTGC |
